# GRAS1 non-coding RNA protects against DNA damage and cell death by binding and stabilizing NKAP

**DOI:** 10.1101/2023.06.20.545783

**Authors:** Tong Su, Nhu Trang, Jonathan Zhu, Lingbo Kong, Darin Cheung, Vita Chou, Lauren Ellis, Calvin Huang, Nichelle Camden, Colleen A. McHugh

## Abstract

Non-coding RNA (ncRNA) gene products are involved in diverse biological processes including splicing, epigenetic regulation, gene expression, proliferation, and metabolism. The biological mechanisms by which ncRNAs contribute to cell survival remain poorly understood. We found that the Growth Regulator Antisense 1 (GRAS1) long non-coding RNA (lncRNA) transcript promotes growth in multiple human cell types by protecting against DNA damage. Knockdown of GRAS1 induced DNA damage and cell death, along with significant expression changes in DNA damage response, intrinsic apoptotic signaling, and cellular response to environmental stimulus genes. Extensive DNA damage occurred after GRAS1 knockdown, with numerous double strand breaks occurring in each cell. The number of cells undergoing apoptosis and with fragmented nuclei increased significantly after GRAS1 knockdown. We used RNA antisense purification and mass spectrometry (RAP-MS) to identify the NF-κB activating protein (NKAP) as a direct protein interaction partner of GRAS1 lncRNA. NKAP protein was degraded after GRAS1 knockdown, in a proteasome-dependent manner. Overexpression of GRAS1 or NKAP mitigated the DNA damage effects of GRAS1 knockdown. In summary, GRAS1 and NKAP directly interact to protect against DNA damage and cell death in multiple human cell lines.

## Introduction

Non-coding RNAs (ncRNAs) are a large class of biomolecules with the potential to regulate cell growth processes. Dysregulated ncRNA expression has been associated with cancer development and metastasis in many cell types (Kazemzadeh et al. 2015, Wang and Chang, 2011). Direct interactions between non-coding RNA and proteins can regulate critical biological processes in humans. For example, XIST RNA coordinates the spatial and temporal localization of the SHARP, N-COR2, and HDAC3 complex to initiate epigenetic silencing across the X chromosome during female development (McHugh et al. 2015). Most non-coding RNA functions remain unknown in human cells. Here, we examined the mechanism of the Growth Regulator Antisense 1 (GRAS1) long non-coding RNA (lncRNA) in promoting human cancer cell growth.

GRAS1 is a lncRNA produced by the EPB41L4A-AS1 gene locus, which was previously identified as a growth regulator in a CRISPR/Cas9 screen (Liu et al. 2017). Altered expression from the gene locus EPB41L4A-AS1 affects cell survival across a broad range of cancer types including gastric (Delshad et al. 2019), lung (Wang et al. 2020), colorectal (Bin et al. 2021), cervical (Liao et al. 2019), placental villus (Zhu et al. 2019), and bone marrow-derived mesenchymal stem cells (Cui et al. 2020). Based on these previous studies, we postulated that GRAS1 lncRNA plays a protective role against cell death in multiple cancer cell types through a common mechanism.

NF-κB activating protein (NKAP) is a multifunctional protein which promotes growth and development in highly proliferative human cells (Pajerowski et al. 2009, Shapiro et al. 2019, Okuda et al. 2015). NKAP was first identified in yeast (Chen et al. 2003) and has since been characterized as an RNA-binding protein in humans (Burgute et al. 2014). NKAP dysregulation is associated with gastric cancer (Wei et al. 2019), colon cancer (Shu et al. 2019), glioblastoma (Sun et al. 2022) and renal cell carcinoma (Ma et al. 2020). No known deletions or truncation mutants of NKAP have been identified in humans, suggesting that it is an essential protein.

Mutations in the C-terminal region of NKAP have been implicated as the cause of human disease in several children displaying neurological and developmental delays (Fiordaliso et al. 2019).

These patients had clinical phenotypes consistent with Hackmann-Di Donato-type X-linked syndromic intellectual disability, a disease which was previously linked to NKAP mutation (Hackmann et al., 2015). Here, we used a combination of molecular biology and biochemistry techniques to uncover a direct interaction between GRAS1 RNA and NKAP. We further identified the mechanism of action of the non-coding RNA GRAS1 in promoting human cell survival through binding to the NKAP protein and preventing its degradation.

## Results

### GRAS1 RNA promotes cell survival in multiple cell types

The GRAS1 long non-coding RNA is located at Chromosome 5q22.1 in the EPB41L4A-AS1 locus, which is associated with the NCBI RefSeq transcript NR_015370.2. Examination of RNA expression in ENCODE RNA-seq data from this locus revealed that an alternative unannotated RNA isoform, termed GRAS1 (Transcript coordinates, hg38 Chr5: 112,160,901-112,162,501), is predominantly expressed in many cell types from this gene locus (Figure 1A). An analysis of RNA expression using The Cancer Genome PanCancer Atlas showed that GRAS1 expression is frequently dysregulated in multiple cancer types, further supporting a potential role for GRAS1 in cell survival (Supplementary Fig. 1) (Cerami et al. 2012, Gao et al. 2013). The subcellular localization of GRAS1 lncRNA was evaluated in A549 lung cancer cells, and GRAS1 was detected in both nuclear and cytoplasmic fractions by qRT-PCR. Control GAPDH mRNA was localized predominantly to the cytoplasm and U1 snRNA was localized predominantly to the nucleus, in line with the known cellular functions of these RNA transcripts (Supplementary Fig. 2A). Based on this dual localization, GRAS1 may function in the nucleus and/or the cytoplasm.

**Figure 1:**
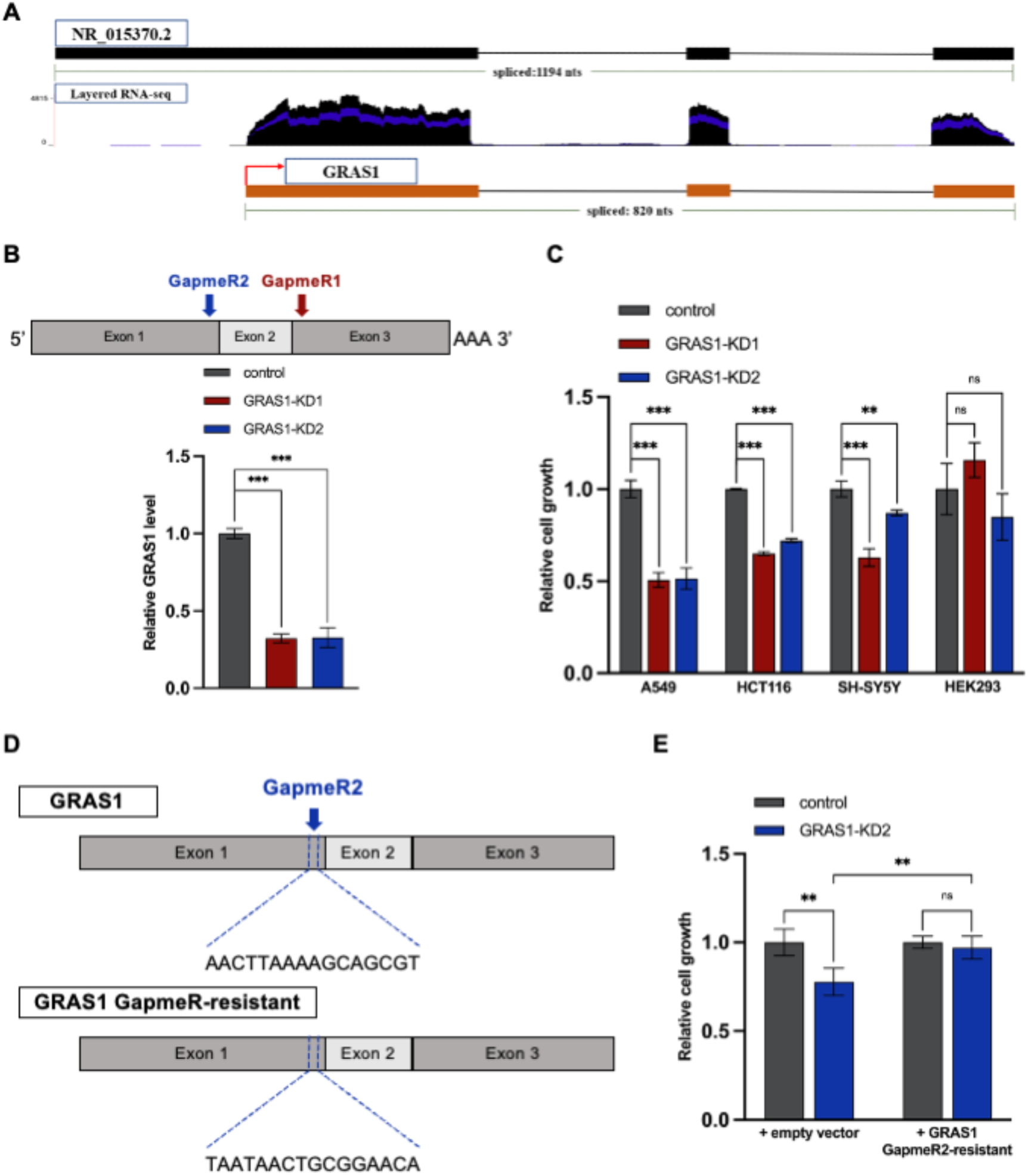
GRAS1 RNA promotes cell survival in multiple cell types. (A) GRAS1 long non-coding RNA locus and ENCODE RNA-seq expression data from the UCSC Genome Browser. (B) Analysis of GRAS1 knockdown efficiency in A549 cells transfected with control GapmeR compared with GRAS1 GapmeR1 or GapmeR2, normalized to GAPDH control mRNA. (C) CCK8 assay for survival of A549, HCT116, SH-S5Y5, and HEK293 cells transfected with control GapmeR compared with GRAS1 GapmeR1 or GRAS1 GapmeR2. (D) Schematic gene structure of GRAS1 GapmeR-resistant construct. (E) CCK8 assay for survival of A549 GRAS1 knockdown cells complemented with GRAS1 GapmeR-resistant compared with empty vector control (right panel). Error bars represent SD. (ns) P > 0.05; (*) P ≤ 0.05; (**) P ≤ 0.01; (***) P < 0.001 by two-tailed Student’s t-test.

We evaluated whether altering GRAS1 expression level affects cell survival in one or more human cell types. Since this gene locus was previously independently identified in prior CRISPR/Cas9 screens as a growth-promoting locus, we tested the survival of human cells after knockdown of GRAS1 with two specific antisense oligonucleotides(ASO) locked nucleic acid GapmeRs. The ASO GapmeR knockdown method of GRAS1 manipulation was chosen, rather than deletion or mutation by CRISPR/Cas9 or other genetic alterations, to avoid changes to the DNA sequence that could affect the function of nearby enhancers or alter the transcriptional expression of neighboring genes. ASO GapmeR targeting of RNA transcripts is highly specific, has high bioavailability, and can achieve RNA target depletion in the nucleus as well as the cytoplasm (Maranon and Wilusz, 2020). We designed two independent ASO GapmeRs targeting different exons of the GRAS1 RNA transcript, and which did not show significant off-target binding to other transcripts in the human genome. A reduction of around 60% of the GRAS1 RNA level was reproducibly achieved after treatment with either of the GRAS1-targeting ASO GapmeRs, compared to the non-targeting control GapmeR (Fig. 1B). Knockdown of GRAS1 with either ASO GapmeR dramatically decreased the survival of A549 non-small cell lung cancer cells, HCT116 colon carcinoma cells, and SH-SY5Y neuroblastoma cells, as measured by mitochondrial function assay, while HEK293 human embryonic kidney cells remained unaffected (Fig. 1C). In addition, colony formation assays confirmed that GRAS1 knockdown with GapmeR1 or GapmeR2 significantly impaired cell growth in A549 lung cancer cells (Supplementary Fig. 3). To further validate the observed phenotypes, GRAS1 knockdown cells were complemented by expression of a GRAS1 GapmeR-resistant construct, with the GapmeR targeting sequence shuffled to prevent degradation by ASO GapmeR (Fig. 1D). The impact of GRAS1 depletion on cell viability was rescued by the complementation of GRAS1 construct, confirming that GRAS1 lncRNA is essential for cell growth and survival (Fig. 1E).

### GRAS1 knockdown results in upregulation of DNA damage response genes

Since knockdown of GRAS1 led to a dramatic decrease in cell survival, we analyzed the differentially expressed genes after GRAS1 knockdown in A549 lung cancer cells to identify biological pathways that were potentially affected by this transcript and could contribute to the observed cell death effects. Transcriptome-wide changes in RNA expression in GRAS1 knockdown cells were measured by Illumina sequencing after 48 hours of GapmeR treatment, with between 2 x 10^7^ and 4 x 10^7^ reads per sample (Supplementary Fig. 4A). Principal Component Analysis (PCA) showed clustering of the three independent biological replicates for each treatment, with a clear separation between knockdown and control samples (Fig. 2A). The knockdown efficiency of GRAS1 in the treated samples was confirmed using normalized read counts from the aligned RNA-seq data. On the contrary, the expression levels of the neighboring antisense gene EPB41L4A and a snoRNA contained within an intron of the GRAS1 transcript were not significantly affected by the ASO GapmeR treatment (Supplementary Fig. 4B). In addition, GRAS1 knockdown did not change the expression level of the overlapping TIGA1 protein in either A549 or HCT116 cells, as evidenced by Western blotting (Supplementary Fig. 4C and D). We confirmed the specificity of GRAS1 ASO GapmeR knockdown for the mature GRAS1 RNA transcript and not the neighboring genes.

**Figure 2:**
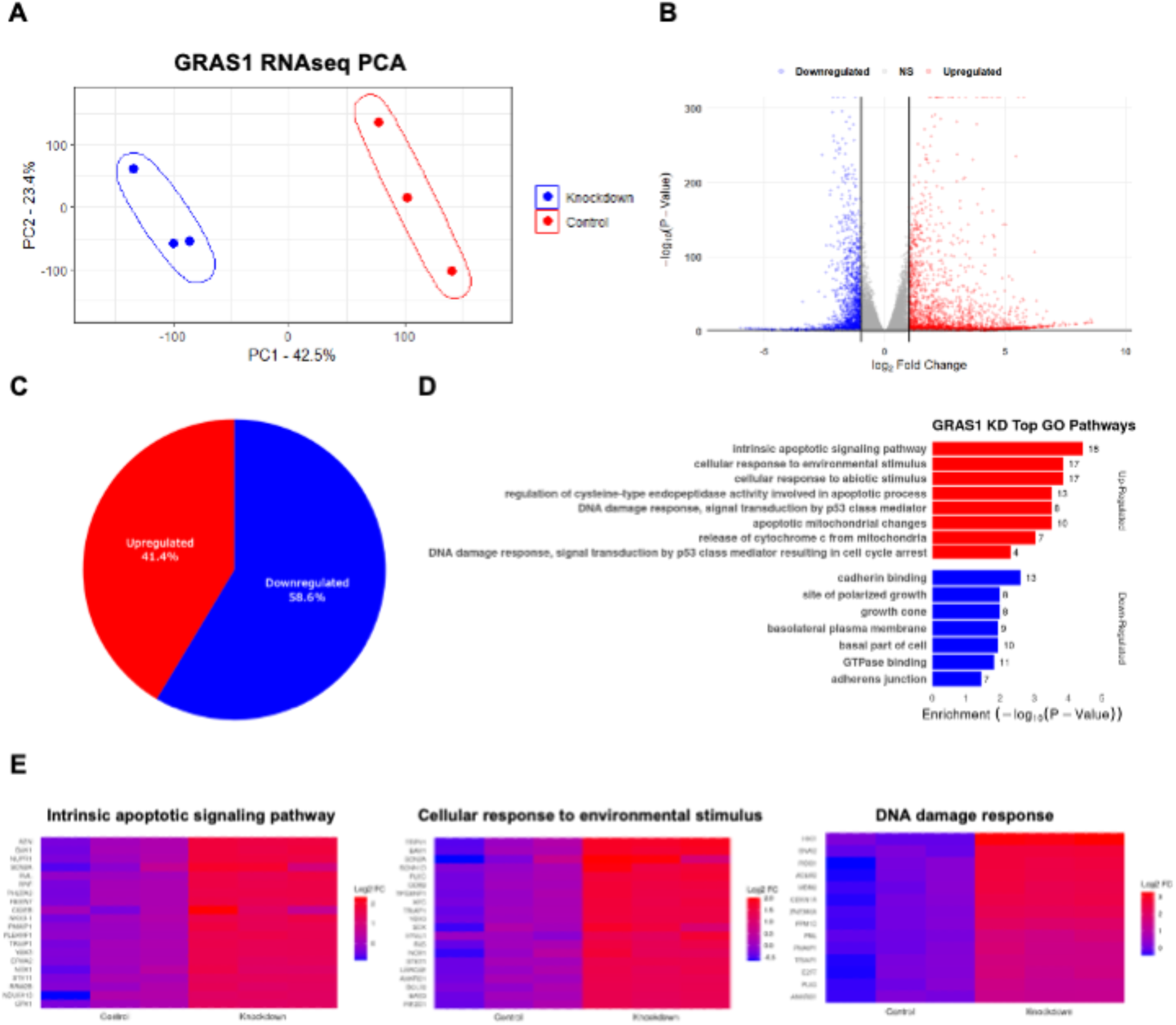
GRAS1 knockdown results in upregulation of DNA damage response genes. (A) Principal component analysis (PCA) of normalized RNA-seq transcripts per million (TPM) from RNA sequencing of triplicate biological samples of A549 cells transfected with control GapmeR (red) compared with GRAS1 GapmeR1 (blue). (B) Volcano plot of differentially expressed genes in control GapmeR compared to GRAS1 GapmeR1 knockdown samples in A549 RNA-seq data. Red and blue points mark the genes with significantly increased or decreased expression, respectively. (C) Percentage of upregulated (red) and downregulated (blue) differentially expressed genes in GRAS1 knockdown cells compared to control cells. (D) Gene Ontology enrichment analysis of selected biological processes in up- and down-regulated genes between control GapmeR and GRAS1 GapmeR1 transfected A549 cells. (E) Relative expression of the top genes from the intrinsic apoptotic signaling pathway, the cellular response to environmental stimulus, and the DNA damage response after GRAS1 knockdown.

Knockdown of GRAS1 non-coding RNA led to significant changes in RNA expression in A549 lung cancer cells. A total of 12,062 genes were more than 2-fold differentially expressed with *p* value ≤ 0.05 (Fig. 2B). In the differentially expressed gene set after GRAS1 knockdown, 41.4% of RNA transcripts were upregulated and 58.6% of RNA transcripts were downregulated (Fig. 2C). The top 1,000 differentially expressed genes based on *p* value from expression level changes in RNA-seq from triplicate biological samples of GRAS1 knockdown versus control GapmeR treatment were used for gene ontology (GO) enrichment analysis. GO term analysis of the differentially expressed genes in GRAS1 knockdown cells identified a significant increase in multiple biological pathways related to cell survival and DNA damage response, and a significant decrease in genes related to polarized cell growth (Fig. 2D). The strongly upregulated transcripts in GRAS1 knockdown cells clustered in pathways including intrinsic apoptotic signaling, cellular response to environmental stimulus, and DNA damage response (Fig. 2E). We found that GRAS1 knockdown leads to differential expression of many mRNA transcript levels, with a significant increase in transcripts associated with maintenance of genome integrity through the DNA damage response.

### GRAS1 knockdown causes accumulation of multiple DNA damage foci in nuclei

Based on the gene expression alterations in pathways related to apoptotic signaling, cellular response, and DNA damage response, we next investigated whether extensive DNA damage occurred in GRAS1 knockdown cells. The TP53 (p53) protein is a key regulator of cell viability and cell cycle progression in response to DNA damage and other types of cell stress (Kastan et al. 1995). We validated that knockdown of GRAS1 significantly increased p53 protein levels (Fig. 3A). Consistent with these results, knockdown of GRAS1 also increased the mRNA expression of p53 downstream gene transcriptional targets including PML, PLK3, and Gadd45a (Fig. 3B). In conclusion, we found that the downregulation of GRAS1 lncRNA transcript expression activates p53 signaling in A549 cells.

**Figure 3:**
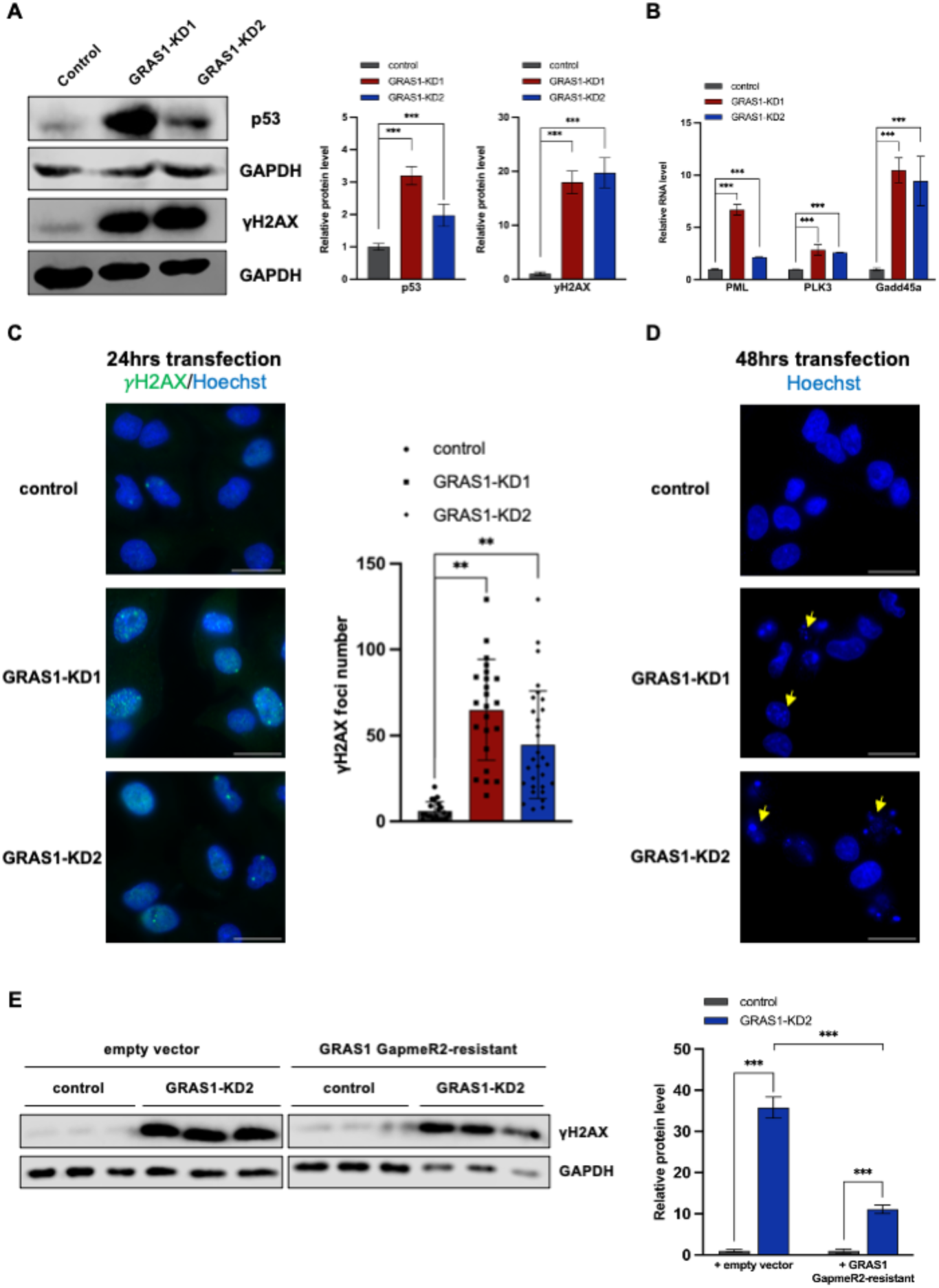
GRAS1 knockdown causes accumulation of multiple DNA damage foci in nuclei. (A) Western blot of p53 and γH2AX in A549 cells treated with control GapmeR or GRAS1 GapmeRs (left panel). Quantification of p53 and γH2AX expression was calculated after normalization to the GAPDH loading control (right panel). (B) Expression of PML, PLK3, and Gadd45a mRNA levels in A549 cells treated with control GapmeR compared with GRAS1 GapmeRs by RT-qPCR. (C) Measurement of γH2AX foci (green) and Hoechst DNA stain (blue) in A549 nuclei after 24 hours of transfection with GRAS1 GapmeRs compared to control GapmeR. Representative images (left panel) and the quantification of γH2AX foci per cell from 3 independent experiments (right panel). Scale bars, 25 μm. (D) Representative images of Hoechst DNA stain (blue) in A549 cells treated with control GapmeR compared with GRAS1 GapmeR1 or GRAS1 GapmeR2 after 48 hours. Yellow arrows indicate nuclear fragmentation. Scale bars, 25 μm. (E) Western blot of γH2AX in A549 cells co-transfected with empty vector or GRAS1 GapmeR-resistant vector, and with control GapmeR or GRAS1 GapmeR2 (left panel). Quantification of γH2AX after normalization to the GAPDH loading control (right panel). Error bars represent SD. (**) P ≤ 0.01; (***) P < 0.001 by two-tailed Student’s t-test.

The p53 signaling pathway can be activated by multiple signals, including DNA damage (Lakin and Jackson, 1999). Based on the observation of a strong upregulation of DNA double-strand break repair genes in the RNA-seq analysis, we next evaluated whether knockdown of GRAS1 increases DNA damage. The histone variant H2AX is rapidly phosphorylated at Serine 139 (γH2AX) in an early cellular response to DNA double strand breaks (Mah et al. 2010). Histone H2AX phosphorylation was measured by Western blot in GRAS1 knockdown cells as an indicator of DNA damage. Knockdown of GRAS1 significantly increased the overall levels of γH2AX in the cell population compared to control cells (Fig. 3A). We then enumerated γH2AX foci in individual cells using immunofluorescence microscopy. Discrete γH2AX foci in cell nuclei correlate with the locations of DNA double-strand breaks (Mah et al. 2010). Consistent with the results of γH2AX immunoblotting, we found that knockdown of GRAS1 significantly increased the number of γH2AX foci, with many individual cells containing more than one hundred foci after 24 hours of GapmeR treatment (Fig. 3C). In comparison, control cells generally showed fewer than ten γH2AX foci per cell. These results indicate that downregulation of GRAS1 leads to a profound increase in DNA damage and double strand breaks in A549 cells. Over time, extensive nuclear fragmentation became apparent in the GRAS1 knockdown treated cells (Fig. 3D). The DNA damage induced by GRAS1 knockdown was significantly reduced by complementation with GRAS1 GapmeR-resistant overexpression, confirming the direct role of GRAS1 in protecting cells against DNA damage (Fig. 3E).

Chromosome condensation and DNA fragmentation is a typical characteristic of programmed cell death after extensive DNA damage (Toné et al. 2007). Knockdown of GRAS1 severely affected the nuclear morphology in A549 cells after 48 hours of GapmeR treatment (Fig. 3D). Regulation of the cell cycle plays a critical role in maintaining cell viability by ensuring that the cell replicates and divides correctly with appropriate DNA content and integrity through each phase of the cycle. To assess the role of GRAS1 in cell cycle regulation, we performed propidium iodide (PI) staining and flow cytometry analysis on GRAS1 knockdown or overexpression cells. Downregulation of GRAS1 led to a significant increase in the number of cells with low DNA content (sub-G1 phase) compared to control cells, indicating that a larger proportion of cells in the population were undergoing apoptosis or contained fragmented chromosomes after GRAS1 knockdown (Supplementary Fig. 5A). Loss of GRAS1 also decreased the proportion of cells in G1 phase. These data indicate that depletion of GRAS1 RNA affects the cell cycle distribution of A549 cells in culture. Cleaved PARP increased in the GRAS1 ASO GapmeR treated cells compared to control cells, indicating cell death due to apoptosis (Supplementary Fig. 5B). Overall, we found that the downregulation of GRAS1 resulted in extensive DNA damage through accumulation of double-strand breaks in A549 cells, leading to cell death by p53 activation and apoptosis.

### GRAS1 non-coding RNA directly binds NF-**κ**B activating protein (NKAP)

To investigate the biological mechanism by which GRAS1 regulates cell growth, we identified direct protein interaction partners of the endogenous GRAS1 RNA transcript in A549 cells using RNA antisense purification with mass spectrometry (RAP-MS). RAP-MS identifies only direct and specific RNA-protein interaction partners by covalently crosslinking “zero-distance” binding interactions between nucleobases and aromatic amino acids with UV254nm light. The use of a unique and highly denaturing wash buffer containing 4 M urea and 500 mM lithium chloride also eliminates indirect and non-specific protein interactions from the capture sample (McHugh et al. 2015, McHugh, Russell and Guttman 2015, Trang et al. 2023). First, the efficient capture and recovery of endogenous GRAS1 RNA with biotinylated DNA antisense probes was validated (Fig. 4A). Endogenous GRAS1 RNA-protein complexes were then purified from UV254nm crosslinked cells to identify direct and specific protein binding partners of the GRAS1 lncRNA. A U1 snRNA capture was used as a positive control and a luciferase mRNA capture was used as a negative control for RAP-MS experiments (Fig. 4B). LC-MS/MS analysis identified 63 proteins in the luciferase control capture, 256 proteins in the U1 control capture, and 192 proteins in the GRAS1 capture sample. After subtracting background proteins that were also identified in the luciferase negative control sample, 138 proteins were specific for the GRAS1 capture sample and 208 proteins were specific for the U1 capture sample. To eliminate commonly identified RNA-binding proteins and splicing factors, we next removed any proteins which were also found in the U1 snRNA capture from the GRAS1 protein list. Finally, the GRAS1 protein list was filtered to include only proteins that were identified with at least two unique peptides, leaving a total of 21 proteins identified as high confidence, direct interaction partners of GRAS1 lncRNA by RAP-MS (Fig. 4C).

**Figure 4:**
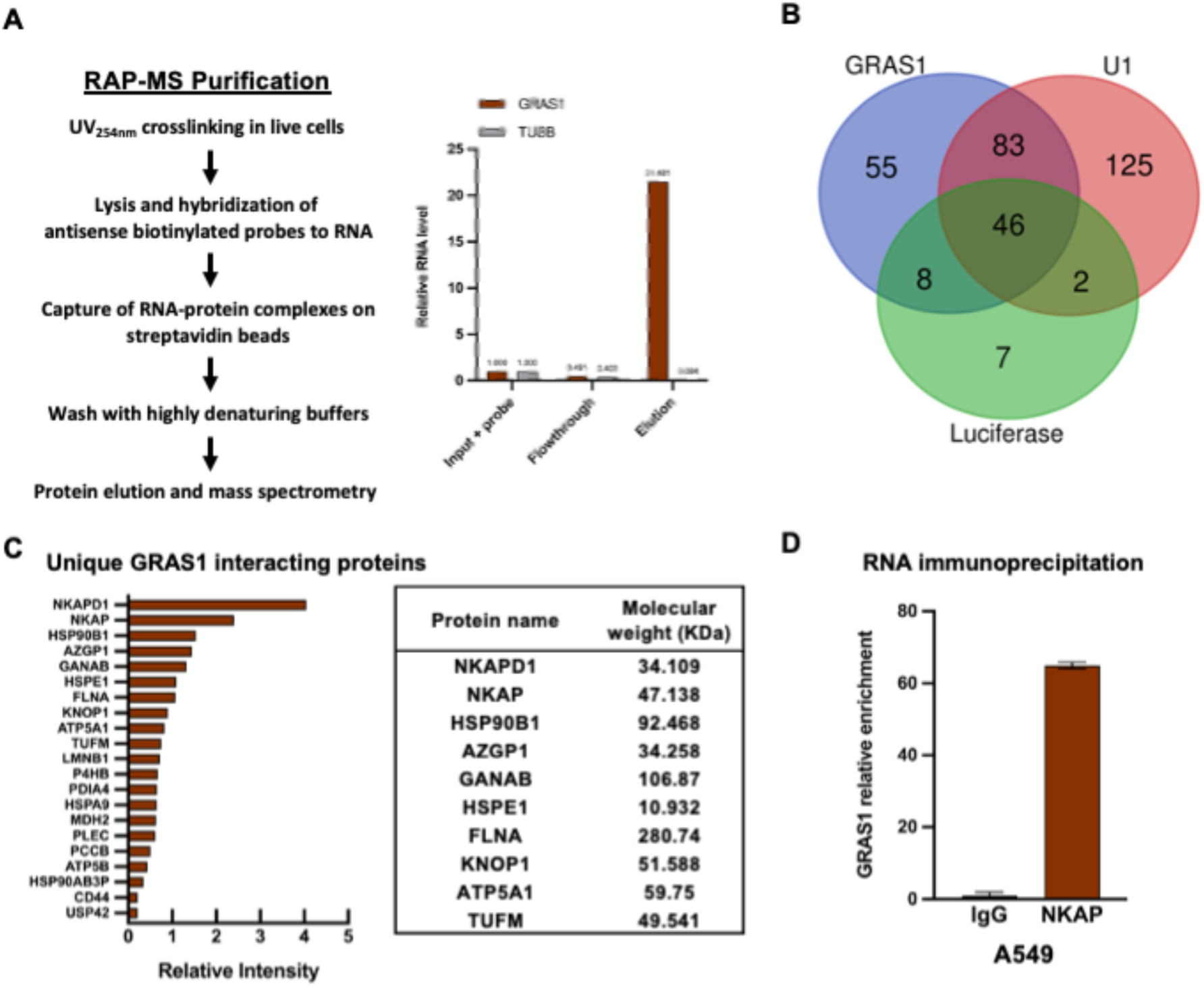
GRAS1 non-coding RNA directly binds NF-κB activating protein (NKAP). (A) RNA antisense purification – mass spectrometry experimental outline (left panel) and the validation of GRAS1 RNA capture and recovery by biotinylated antisense oligonucleotide probes against GRAS1 (right panel). RT-qPCR on GRAS1 RNA and TUBB control mRNA in three fractions, including input with probes added, unbound sample flowthrough, and elution. (B) Total proteins identified in RAP-MS of GRAS1 RNA, U1 snRNA positive control, and luciferase mRNA negative control elution samples. (C) GRAS1 interactome proteins identified by RAP-MS, with at least two unique peptides, after excluding proteins that were also found in U1 and luciferase control captures (left panel). The top ten GRAS1-interacting proteins and their molecular weights (right panel). (D) GRAS1 RNA enrichment measured by RT-qPCR after immunoprecipitation of NKAP in A549 cells, compared to immunoprecipitation with IgG control antibody.

NKAP and C11ORF57/NKAPD1 were identified in our MS screen as the top interaction partners of GRAS1 lncRNA. Consistent with direct binding to GRAS1 RNA, NKAP was previously characterized as an RNA-binding protein (Burgute and Noegel, 2014). To validate the interaction between GRAS1 and NKAP by an orthogonal approach, we performed RNA immunoprecipitation (RIP) in non-crosslinked A549 cell lysates, by capturing NKAP and testing for the presence of GRAS1 by qRT-PCR. We observed more than a 50-fold increase in GRAS1 RNA capture in NKAP-IP sample elution compared to the control IgG sample elution (Fig. 4D). The function of C11ORF57/NKAPD1 remains uncharacterized. Therefore, we focused on the NKAP protein to investigate whether the direct interaction between GRAS1 and NKAP contributes to cell survival and protection against DNA damage.

### GRAS1 stabilizes NKAP to protect against cell death and DNA damage

Since GRAS1 directly interacts with NKAP, we investigated the effect of GRAS1 knockdown on NKAP mRNA and protein levels. Notably, loss of GRAS1 had no significant effect on NKAP mRNA levels (Fig. 5A) but caused a significant decrease in the protein level of NKAP (Fig. 5B). Therefore, we hypothesized that GRAS1 RNA interaction affects NKAP stability through a post-translational mechanism. Treatment of cells with the proteasome inhibitor MG-132 indeed reduced the effect of GRAS1 knockdown on the protein level of NKAP (Fig. 5B). The combined data from RAP-MS and RNA immunoprecipitation experiments indicated that GRAS1 lncRNA binds directly to NKAP in A549 cells, and that GRAS1 RNA binds and protects NKAP from proteasomal degradation through a post-translational mechanism.

**Figure 5:**
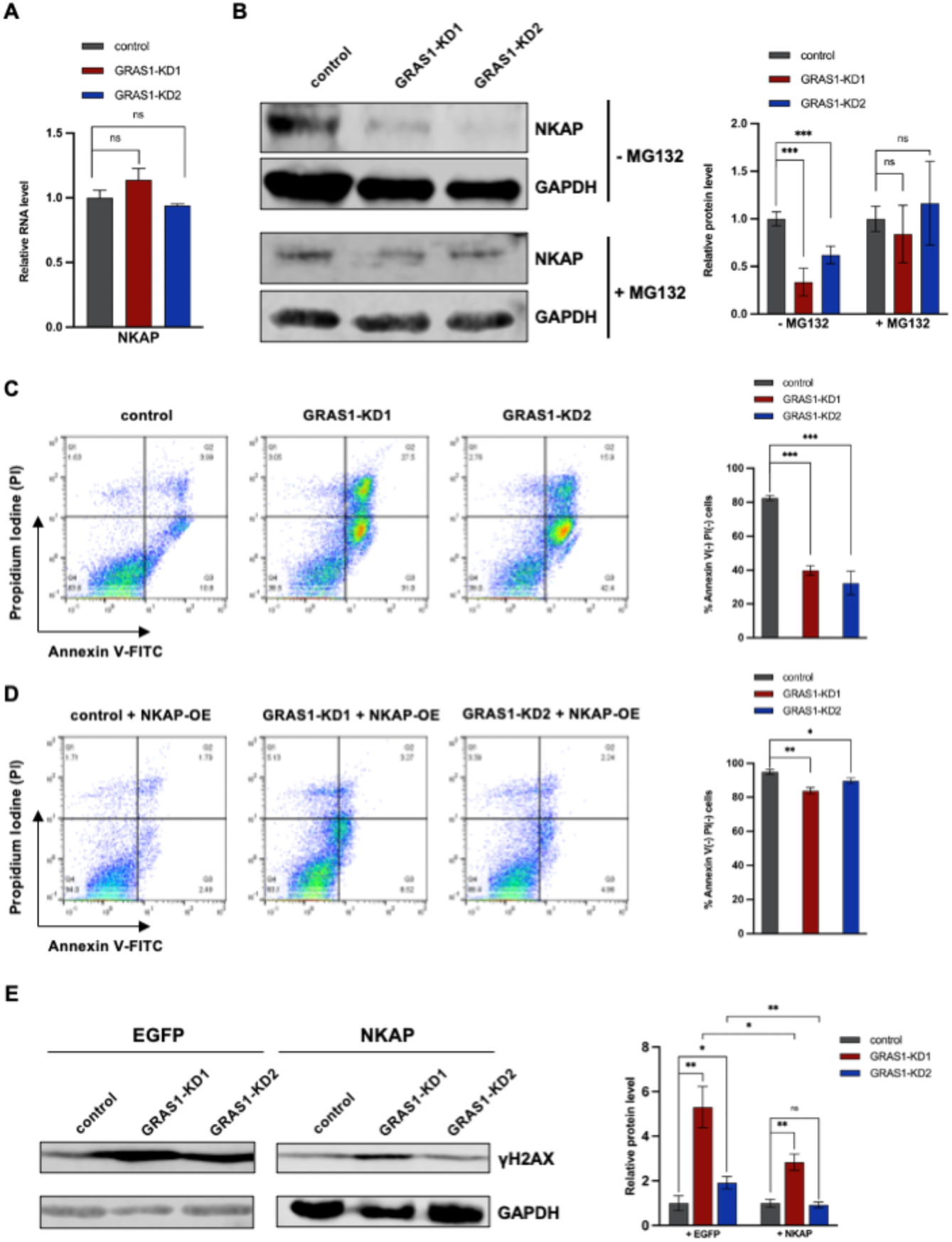
GRAS1 stabilizes NKAP to protect against cell death and DNA damage. (A) RT-qPCR for NKAP mRNA expression in A549 cells with control GapmeR or GRAS1 GapmeRs (B) Western blot of NKAP in control GapmeR treated A549 cells compared to GRAS1 GapmeRs knockdown in standard medium (top) or with 5 µM MG132 treatment (bottom). Quantification of NKAP expression was calculated after normalizing to GAPDH loading control for each sample (right panel). (C) GRAS1 knockdown induced A549 cell death, as assessed by flow cytometry with Annexin V/PI staining. Representative results of A549 with control GapmeR, GRAS1 GapmeR1 or GRAS1 GapmeR2 treatment (left panel). Quantification on percentage of Annexin V-negative/ PI-negative A549 cells treated with control GapmeR compared with GRAS1 GapmeR1 or GRAS1 GapmeR2 (right panel). (D) NKAP overexpression increases the percentage of live cells from GRAS1 knockdown, as assessed by flow cytometry with Annexin V/PI staining. Representative results (left panel), and quantification of cells (right panel). (E) Western blot of γH2AX in A549 cells co-transfected with control EGFP vector or NKAP vector, and with control GapmeR, GRAS1 GapmeR1, or GRAS1 GapmeR2 (left panel). Quantification of γH2AX expression was calculated after normalization to the GAPDH loading control for each sample (right panel). Error bars represent SD. (ns) P > 0.05; (*) P ≤ 0.05; (**) P ≤ 0.01; (***) P < 0.001 by two-tailed Student’s t-test.

To determine whether DNA damage resulting from NKAP depletion is a main contributor to cell death after GRAS1 knockdown, we performed rescue experiments with NKAP expression in GRAS1 knockdown cells. As observed previously, GRAS1 GapmeR treatment significantly decreased cell survival, compared to cells treated with control GapmeR (Fig. 5C). By flow cytometry, the apoptotic cell population marked by Annexin V staining was significantly increased in GRAS1 knockdown cells. We observed that NKAP expression protected cells from apoptotic death and abrogated the effect of GRAS1 knockdown in the Annexin V and PI staining assay (Fig. 5D). Finally, we tested if expression of NKAP could reverse the effect of GRAS1 depletion on DNA damage. Overexpression of NKAP slightly decreased the levels of γH2AX in GRAS1 knockdown cells treated with ASO GapmeR 1 and significantly decreased the γH2AX levels in GRAS1 knockdown cells treated with ASO GapmeR 2 (Fig. 5E). In summary, we found that NKAP overexpression protects against the cellular effects of GRAS1 downregulation.

### GRAS1 function is conserved in HCT116 colorectal cancer cells

Our initial cell proliferation assay screen indicated that GRAS1 is essential for cell growth and survival in multiple cell lines (Fig. 1C). In A549 cells, we found that GRAS1 knockdown resulted in a significant increase in DNA damage and cell death, and that GRAS1 interacted directly with NKAP. We further investigated whether GRAS1 affects similar cellular pathways in HCT116 colorectal cancer cells. We found that GRAS1 knockdown also resulted in a significant increase in γH2AX protein levels in HCT116 cells, demonstrating an induction of DNA damage leading to cell death (Fig. 6A). This effect was reversed by the overexpression of GRAS1 GapmeR-resistant RNA, confirming the direct effect of GRAS1 RNA on HCT116 cell death (Fig. 6B). In addition, we validated the interaction between GRAS1 and NKAP in HCT116 cells by RNA immunoprecipitation assay and observed a profound increase in GRAS1 captured by NKAP compared to the control IgG capture (Fig. 6C). Furthermore, the overexpression of NKAP partially protects cells against DNA damage induced by GRAS1 depletion (Fig. 6D).

**Figure 6:**
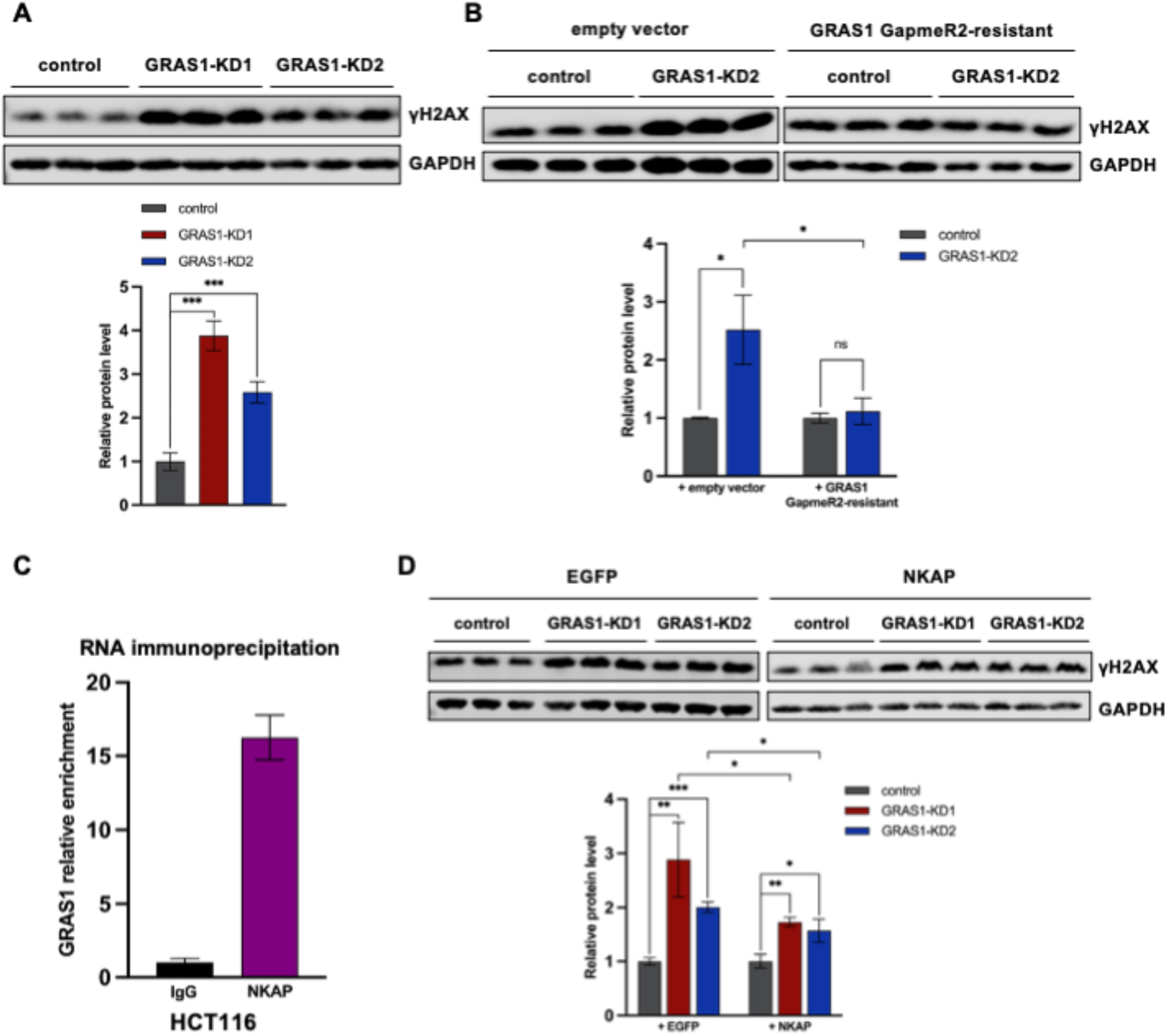
GRAS1 function is conserved in HCT116 colorectal cancer cells. (A) Western blot of γH2AX in HCT116 cells treated with control GapmeR, GRAS1 GapmeR1 or GRAS1 GapmeR2 (top). Quantification of γH2AX expression was calculated after normalization to the GAPDH loading control for each sample (bottom). (B) Western blot of γH2AX in HCT116 cells co-transfected with control empty vector or GRAS1 GapmeR-resistant vector, and with control GapmeR or GRAS1 GapmeR2 (top). Quantification of γH2AX expression was calculated after normalization to the GAPDH loading control for each sample (bottom). (C) GRAS1 RNA enrichment measured by RT-qPCR after immunoprecipitation of NKAP in HCT116 cells, compared to immunoprecipitation with IgG control antibody. (D) Western blot of γH2AX in HCT116 cells co-transfected with control EGFP vector or NKAP vector, and with control GapmeR, GRAS1 GapmeR1, or GRAS1 GapmeR2 (top). Quantification of γH2AX expression was calculated after normalization to the GAPDH loading control for each sample (bottom). Error bars represent SD. (ns) P > 0.05; (*) P ≤ 0.05; (**) P ≤ 0.01; (***) P < 0.001 by two-tailed Student’s t-test.

Overall, we found that GRAS1 function is conserved in HCT116 colorectal cells, confirming that the interaction between GRAS1 and NKAP is important for DNA damage protection in multiple cell types.

## Discussion

Non-coding RNAs have been proposed to function in critical cell survival pathways, but their mechanisms generally remain unknown. We identified a new mechanism for the GRAS1 non-coding RNA in promoting cell survival and preventing DNA damage in multiple human cell lines. Depletion of GRAS1 RNA with ASO GapmeRs led to a significant increase in cell death, while complementation with a GapmeR-resistant version of the GRAS1 RNA, or its directly interacting protein factor NKAP, was protective against cell death and DNA damage in lung and colon cancer cell lines. This is the first report of a specific RNA-protein interaction affecting NKAP function in human cells.

Initial mechanistic studies of GRAS1 RNA function were performed in the non-small cell lung cancer cell line A549, to investigate the phenotypic effects of GRAS1 knockdown and identify the cause of cell death. We found that extensive DNA damage occurred in GRAS1-depleted cels. Transcriptomic analysis of differentially expressed genes in GRAS1 knockdown cells compared to control cells revealed the upregulation of pathways related to cell survival, including p53 activation and DNA damage response. Knockdown of GRAS1 increased p53 protein levels and increased the mRNA levels of p53 target genes in A549 cells. Mammalian cells generally maintain a low level of p53, due to continual degradation and rapid turnover.

Cellular p53 accumulation can be caused by multiple stressors, including DNA damage (Lakin and Jackson 1999). We investigated the protective effect of GRAS1 on genome integrity in A549 cells. GRAS1 knockdown significantly increased the number of double-strand DNA breaks on chromatin, enumerated by staining of γH2AX foci in nuclei. In an individual cell, more than one hundred double-strand breaks were frequently observed after depletion of GRAS1 RNA. GRAS1 depletion also affected progression through the cell cycle in A549 cells. A significant increase in the sub-G1 cell population indicated that GRAS1 knockdown results in an increased number of cells with low DNA content, in line with our microscopic studies of extensive DNA damage and nuclear fragmentation after GRAS1 depletion. The damage induced by GRAS1 knockdown led to apoptotic cell death, as indicated by PARP cleavage and Annexin V flow cytometry.

To identify protein interaction partners of the endogenous GRAS1 RNA, we used an updated version of the UV-crosslinking and a highly denaturing mass spectrometry purification technique, RAP-MS (Trang et al. 2023). RAP-MS identified 21 proteins that directly bind to and uniquely interact with GRAS1, compared to luciferase and U1 snRNA controls. A top GRAS1 binding partner was NKAP, a protein which promotes accurate chromosome segregation and cell survival. The interaction between NKAP RNA and GRAS1 protein was further validated by RNA immunoprecipitation. Knockdown of GRAS1 ncRNA reduced NKAP protein levels, while the expression of NKAP mRNA was unaffected. Inhibition of proteasomal degradation prevented NKAP degradation upon GRAS1 knockdown. Therefore, we concluded that GRAS1 binding affects NKAP stability through a post-translational regulation mechanism.

Next, we investigated the cause of death in GRAS1 knockdown cells. We found that the interaction between GRAS1 and NKAP protects against extensive DNA damage and cell death. The decrease in cell survival after GRAS1 knockdown could be rescued by ectopic expression of NKAP, indicating that GRAS1-NKAP stabilization is directly related to the cell survival in A549 cells. Notably, overexpression of NKAP mitigated the DNA damage effects of GRAS1 knockdown in A549 cells. Dysregulation of NKAP has been linked to chromosome instability and DNA damage in previous work. Depletion of NKAP causes chromosome segregation defects and genome instability in HeLa cells (Li et al. 2016) and increases the number of DNA damage foci in U2-OS osteosarcoma cells (Zhang et al. 2023). Mutations in NKAP have recently been identified as a cause of a neurodevelopmental disorder in multiple unrelated families (Fiordaliso et al., 2019), emphasizing the importance of NKAP function in human cells. Our finding of direct binding to GRAS1 uncovers a new mechanism for regulation of NKAP at the post-translational level. In addition to the initially screened cell lines, we further investigated the functionality of GRAS1 in HCT116 colorectal cancer cells due to the high amplification frequency of GRAS1 in colorectal cancer, which was revealed in analysis of TCGA Pan-Cancer data (Supplemental Figure 1). Our observations of extensive DNA double-strand breaks and death in GRAS1 knockdown cells, along with complementation with the RNA and protein, showed that GRAS1 and NKAP play similar roles in colorectal cancer cells. We concluded that the GRAS1-NKAP interaction protects against DNA damage and promotes cell growth in multiple human cell types.

Cellular functions of many of the non-coding RNAs produced by the human genome are still uncharacterized, and our work advances the current understanding of RNA-protein interactions and their functions in human cells. CRISPR/Cas9 screens have identified growth effects for non-coding RNA transcripts in several cell types, and we investigated the function of one of the broadly expressed RNA hits, the Growth Regulator Specific Antisense 1 (GRAS1) transcript. In our study, GRAS1 RNA promotes cell growth and binds directly to NKAP in multiple cell lines, validating the effect observed in CRISPR/Cas9 screening. Loss of GRAS1 RNA led to DNA damage and cell death, which was be rescued by GRAS1 complementation or overexpression of the directly interacting protein NKAP. We conclude that GRAS1 RNA contributes to the maintenance of cell survival through a direct interaction with the NKAP protein. Post-translational regulation of protein expression levels is an important mechanism of cellular control during cancer development and metastasis. The identification of GRAS1 RNA-mediated regulation of NKAP stability, and the role of the GRAS1-NKAP interaction in protecting multiple cell types against DNA damage and cell cycle arrest, have uncovered a new potential pathway for the manipulation of cancer cell growth control.

## Materials and Methods

### Cell lines and culture conditions

Human lung adenocarcinoma A549 cell line (ATCC No. CCL-185), human colorectal carcinoma HCT116 cell line (ATCC No. CCL-247), human kidney embryonic HEK293 cell line (ATCC No. CRL-1573), and human neuroblastoma SH-SY5Y cell line (ATCC CRL-2266) were purchased from ATCC. A549 and HEK293 cells were cultured in Eagle’s Minimal Essential Medium (EMEM) (Quality Biological, USA); HCT116 cells were cultured in McCoy’s 5A Modified Medium (Gibco, Life Technologies, USA); SH-SY5Y cells were cultured in 1:1 DMEM/F12 media (Gibco, Life Technologies, USA). All mediums were supplemented with 10% fetal bovine serum (Gibco, Life Technologies, USA) and 1% additional L-Glutamine (Corning, USA). All cells were grown in a cell culture incubator at 37 °C with 5% CO2. Cells were tested monthly for mycoplasma.

### Plasmid generation and transfection

Gene inserts were generated by PCR from A549 or K562 complementary DNA (cDNA) using locus-specific primers. Entry vectors of the Gateway cloning system were created with the pENTR/D-TOPO Cloning Kit (2402649, Invitrogen, CA, USA). Once the entry vectors were generated, the gene inserts were transferred to pcDNA3.2 destination vectors using the Gateway LR Clonase Enzyme Mix (11791011, ThermoFisher Scientific, USA). For plasmid transfections, cells were seeded with 2 x 10^5^ cells/well in 12-well plates overnight before each treatment. Cells in each well were transfected with 1 μg of plasmid using 3.5 μL of Lipofectamine 2000 (ThermoFisher Scientific, USA) in Opti-MEM I reduced serum medium (31985062, Life Technologies, USA) following the manufacturer’s suggested protocol.

### Antisense oligonucleotide (ASO) GapmeR knockdown of GRAS1

For GRAS1 RNA knockdown, cells in each well were transfected with 50 μM of each ASO locked nucleic acid GapmeR1 or GapmeR2 (Qiagen, custom design) with Negative Control GapmeR A (30300019-2, Qiagen) using 3.5 μL of Lipofectamine 2000 in Opti-MEM I reduced serum medium following the manufacturer’s suggested protocol. Samples were incubated for 24 or 48 hours before collection, as indicated in the text. GapmeR target sequences were: GRAS1 GapmeR1 sequence (AAATGATCCAGGCAAC) and GRAS1 GapmeR2 sequence (ACGCTGCTTTTAAGTT).

### Nucleic acid isolation and qRT-PCR

Total RNA from all cell treatments was isolated using QIAshredder and RNeasy Mini kit (Qiagen) then further purified or concentrated as needed with RNA Clean & Concentrator kit (Zymo Research). For cell fractionation experiments, total RNA from each cell fraction was isolated using Qiagen RNeasy columns. DNA isolations were performed using plasmid Miniprep kit (Qiagen). All isolations were performed according to the manufacturer’s protocol. 1.5 μg of purified RNA was used for each reverse transcription reaction with 1 μL of Superscript III Reverse Transcriptase (18080085, Invitrogen, USA). Quantitative real-time PCR (qRT-PCR) was performed using ROX Reference Dye (Invitrogen, USA) and SYBR Green Dye (Life Technologies, USA), with fold expression calculated using the 2^-ΔΔCt^ method. Samples were repeated in biological triplicate. Oligonucleotide primers are listed in Supplementary Table 1. Data were normalized to IgG for immunoprecipitation experiments, normalized to GAPDH or Tubulin control genes for qRT-PCR experiments, and normalized to respective RNA amounts for cellular fractionation products.

### Western blotting

Following transfection, cells were harvested with trypsin and washed with 1X PBS. Cells were then mixed with cell lysate buffer, protein inhibitor cocktail (539134, Millipore Sigma), sodium fluoride, and incubated at 95 °C for 10 min before SDS-PAGE separation. Cell lysates were loaded into a 10-well 10% Bis-Tris SDS-polyacrylamide gel along with PageRuler Plus Protein ladder (26619, ThermoFisher Scientific). Proteins were separated by electrophoresis at 130V for 90 min and subsequently transferred to 0.2 micron pore size nitrocellulose membrane.

Membranes were blocked with the LI-COR Biosciences Intercept Blocking buffer (VWR International) for one hour at room temperature and incubated with primary antibodies overnight at 4 °C on a rocker. The following primary antibodies were used: NKAP (ab229096, Abcam), **γ**H2AX (60566SF, Cell Signaling Technology), p53 (10442-1-AP, Proteintech), GAPDH (60004-1-Ig, Proteintech), Lamin A/C (10298-1-AP, Proteintech), PARP1 (13371-1-AP, Proteintech), and TIGA1 (MBS9610109, MyBioSource). All antibodies were diluted in 1X TBST. After incubation, membranes were washed four times for 5 min with 1X TBST buffer (1X TBS with 0.1% Tween-20). Nitrocellulose membrane was then incubated with goat Anti-mouse (102971-154, VWR International) and goat Anti-rabbit (103011-498, VWR International) IRDye-conjugated secondary antibodies for one hour under dark conditions on a rocker. After four 5-min washes, the membrane was imaged with an Odyssey Fc device (LI-COR Bioscience, USA) and analyzed with Empiria Studio software (LI-COR Bioscience, USA).

### Cellular Fractionation

For separation of cytoplasmic and nuclear components, the protocol was performed essentially as previously reported (Gagnon et al. 1999) Briefly, 1 x 10^7^ A549 cells were collected by trypsinization and resuspended in 1 mL hypotonic lysis buffer (10 mM Tris-HCl [pH 7.5], 10 mM NaCl, 3 mM MgCl2, 0.3% NP-40 supplemented with Complete protease inhibitor cocktail (Roche) and 10% glycerol). After 10 min of incubation on ice and a brief vortex of the sample, 60 μL input control for Western Blot was reserved. The remaining lysate was centrifuged for 8 min, 800 x *g* at 4 ℃. The supernatant was separated and saved as the cytoplasmic fraction, and 5 M NaCl was added to create a final concentration of 140 mM NaCl in the sample. The remaining pellet was washed four times by resuspending in hypotonic lysis buffer and centrifuged for 2 min at 300 x *g*. The supernatants were saved as individual washes. Total nuclear extract was then prepared by suspending the pellet in 0.5 mL nuclear lysis buffer (20 mM Tris-HCl [pH 7.5], 150 mM KCl, 3 mM MgCl2, 0.3% NP-40 supplemented with 1 complete protease inhibitor cocktail). The input sample was sonicated in a water bath for 15 min while the nuclear fraction and the four washes were sonicated on ice with microtip at 25% duty and 4 output for 60 seconds each on a Branson ultrasonicator. After sonication, all samples except input were centrifuged at 18,000 x *g* for 10 min at 4 ℃. Supernatants were transferred to fresh tubes and used for Western Blot and RNA isolation. U1 RNA and Lamin A protein were measured as references for the nuclear fraction and GAPDH mRNA and GAPDH protein were measured as references for the cytoplasmic fraction.

### Colony formation assay

For colony formation assays, 2 x 10^5^ A549 cells were seeded in a 6-well plate and incubated overnight, followed by GapmeR knockdown targeting GRAS1. After 24-hour incubation at 37°C and 5% CO2, transfected cells were collected using trypsin (Corning, USA) and resuspended in 4 mL EMEM medium. Then, 50 μL of resuspended cell solution was added to 36.5 cm^2^ cell dishes with 4 mL EMEM medium and incubated for 10 days. Finally, the medium was discarded, and the cells were fixed using 500 μL of fixation buffer (3:1 ratio of methanol to acetic acid). Fixed cells were stained with 0.1% crystal violet (Ricca Chemical, TX, USA).

### Immunofluorescence assay and microscopy

1 x 10^5^ cells were seeded in a tissue culture dish with glass bottom (20102018, FluoroDish, FL, USA) and incubated overnight before transfection. Transfections were performed as described above. After transfection, cells were washed three times with ice-cold 1X PBS, then fixed with chilled 100% methanol at room temperature for 5 min, and washed three times with 1X PBS for 5 min. Cells were permeabilized by treatment with 0.2% Triton X-100 for 10 min at room temperature, then washed three times with 1X PBS for 5 min. Then, cells were incubated in 1% BSA (bovine serum albumin) in 1X PBST (1X PBS + 0.1% Tween 20) for 30 min blocking using Normal Goat Serum (ab7481, Abcam) at room temperature. Cells were incubated in primary antibodies for 1 hour at room temperature and washed three times with 1X PBS. Then cells were incubated with Goat Anti-Rabbit IgG H&L Secondary Antibody (ab6717, Abcam) for 1 hour and washed 3 times with 1X PBS. Finally, cells were incubated in either 1 μg/mL DAPI DNA stain (4083S, Cell Signaling Technology) or 0.5 μg/mL Hoechst 33342 Reagent (R37165, ThermoFisher Scientific, USA) for 1 min, rinsed with 1X PBS, and imaged using the Eclipse Ti Inverted Microscope (Nikon, Japan) with Nikon’s NIS-Elements imaging software.

### Protein immunoprecipitation

For each reaction, 3.5 x 10^7^ A549 cells were harvested and resuspended in 1X PBS, then an equal volume of cold polysome lysis buffer was added (20 mM Tris-HCl [pH 7.5], 50 mM KCl, 10 mM MgCl2, 1 mM DTT, 1X Complete protease inhibitor cocktail (Roche), 200 units/mL RNase inhibitor murine (New England Biolabs)). Cells were lysed by incubation at 4 ℃ for 5 min followed by incubation at -80 ℃ for about two hours before centrifuging for 10 min at 20,000 x *g*. The supernatants were collected and divided so that there were 1 x 10^7^ A549 cells per sample and 5 x 10^6^ cells were saved as whole cell lysate inputs. 75 μL/sample of Dynabeads Protein G (Thermo Fisher Scientific) were washed twice with NT-2 buffer (50 mM Tris-HCl [pH7.5], 150 mM NaCl, 1 mM MgCl2, 0.05% NP-40) and resuspended in 100 μL/sample of NT-2 buffer, with 3 μg of each antibody per sample: IgG as a negative control (10284-1-AP, Proteintech), TDP-43 as a positive control (10782-2-AP, Proteintech), and NKAP. The beads/antibody mixtures were incubated for 90 min at room temperature, then beads were washed six times with NT-2 buffer before resuspending in NET-2 buffer (50 mM Tris-HCl [pH 7.5], 150 mM NaCl, 1 mM MgCl2, 0.05% NP-40, 20 mM EDTA [pH 8.0], 1 mM DTT, 200 units/mL RNase inhibitor murine). Each antibody had the cell lysate supernatant added so the total volume including the NET-2 buffer would be 1 mL. All three samples were incubated overnight at 4 ℃ with gentle rotation, followed by six washes with NT-2 buffer, then each sample was split in half for protein and RNA analysis. For RNA samples, the beads were resuspended in NLS elution buffer (20 mM Tris–HCl [pH 8.0], 10 mM EDTA [pH 8.0], 2% NLS, and 2.5 mM TCEP) for proteinase K digest, then the beads were separated and standard RNA silane cleanup with Dynabeads MyOne Silane (Thermo Fisher Scientific) was performed on the supernatant before reverse transcribing the RNA using SuperScript III to perform qRT-PCR. For protein samples, elution 1 was prepared by using 10 μL of 2X Tris-Glycine SDS sample buffer (Thermo Fisher Scientific) was to resuspend beads containing immunocomplexes and incubated at 50 ℃ for 10 min, then 9 μL of UltraPure water (Invitrogen) and 1 μL of 0.1 M DTT was added to the elution sample. Elution 2 was collected by resuspending the beads in 10 μL of 2X Tris-Glycine SDS sample buffer and 1 μL of 0.1 M DTT and incubating at room temperature for 5 min before adding 9 μL of UltraPure water to bring the final volume of the sample to 20 μL. All protein samples were heated at 95 ℃ for 10 min before storing at -80 ℃.

### Cell proliferation assays

The viability of A549, HCT116, SH-SY5Y and HEK293 cells were determined by modified microculture tetrazolium (MTT) assay (University at Buffalo, State University of New York, NY, USA) or CCK-8 assay (Abcam). Before any treatment, 1.5 x 10^4^ cells were seeded into each well of 96-well flat-bottomed microplates containing medium overnight. The cells were treated and incubated in a humidified incubator in 5% CO2 for 48 hours at 37°C. For MTT assay, growth medium was replaced with the working MTT solution following the manufacturer’s instructions. The resulting solution was incubated in 5% CO2 for another 20 min at 37 °C. Formazan crystals were dissolved in 100 μL of DMSO. For CCK-8 assay, 10 μL WST-8 solution was added to the growth media, and cells were incubated in 5% CO2 for 2 hours at 37 °C. The plates were then analyzed on the Synergy HTX Multi-Mode Microplate Reader (Agilent, USA) at 570 nm. Each sample was assayed with at least three biological replicates.

### Cell cycle analysis with flow cytometry

Following treatment, A549 cells were harvested with trypsin and washed twice with 1X PBS. Cells were then permeabilized and fixed with 70% ice-cold ethanol drop-wise and incubated for 15 min on ice, gently vortexing to prevent clumping. Cells were washed twice with 1X PBS. Then, cells were resuspended in PI Staining Solution (3.8 mM sodium citrate, 50 μg/mL PI (Biotium, USA) and 40 μg/mL RNase A (Zymo Research, USA)) and incubated for 40 min at 37 °C in the dark. Cells were centrifuged and the supernatant was gently aspirated. Finally, the cells were resuspended in 500 μL ice cold 1X PBS and examined on the S3e Cell Sorter (Bio-Rad Laboratories). Cell cycle profiles were analyzed using FlowJo software (BD Biosciences).

### Illumina RNA sequencing

RNA from GRAS1 knockdown in A549 cells was isolated with RNeasy Mini kit and purified using RNA Clean & Concentrator kit. Sequencing was performed for three biological replicates each for both the knockdown and the control samples. Illumina RNA-seq libraries were prepared at the Institution for Genomic Medicine (IGM) using Illumina stranded mRNA prep after poly-A selection. Quality of the libraries was examined by Agilent BioAnalyzer, and sequencing was performed on a NovaSeq S4 with the run type of PE100. Raw paired-end FASTQ sequencing reads were mapped with STAR version 2.7.9a (Dobin et al. 2013) to human reference genome assembly GRCh38 (Schneider et al. 2017) and GENECODE gene annotation file Ensembl 104 (Frankish et al. 2018). FeatureCounts version 2.0.3 (Liao et al. 2014) was used to generate read count matrices, and DESeq2 version 1.34.0 (Love et al. 2014) analysis identified differential expression of RNA transcripts between the triplicate samples of control GapmeR or GRAS1 knockdown GapmeR treated cells. Gene ontology enrichment analysis was performed with enrichGO version 4.2.2 (Yu et al. 2012). RNA sequencing data are available on the NCBI Sequence Read Archive associated with the BioProject PRJNA979253.

### RNA antisense purification with mass spectrometry (RAP-MS)

RAP-MS experiments were performed from A549 cells grown in culture. The method was performed as previously described (Trang et al. 2023) with the following modifications. Cells were collected and crosslinked with 254 nm wavelength light using an XL-1500 Spectrolinker (Spectro-UV, USA). Per 20 M cells, the cell pellet was resuspended in 900 μL cold lysis buffer (10 mM Tris-HCl pH7.5, 500 mM LiCl, 0.5% n-dodecyl-β-D-maltoside, 0.2% sodium dodecyl sulfate, 0.1% sodium deoxycholate) with 1X protease inhibitor cocktail and 920 U of murine RNase inhibitor (M0314L, New England Biolabs, UK). Resuspended cell pellets were further lysed on ice with ten passes through an 18G blunt needle, followed by two sets of 1.5 min sonication with a Branson sonicator using the microtip (setting: 35 duty, 3.0 power output). Cell lysate was incubated with 20 U of TurboDNase (Invitrogen, USA) for 15 min at 37 °C to remove DNA in the lysate, and then quenched by adding 10 mM EDTA, 5 mM EGTA, and 2.5 mM TCEP. The 6 M urea hybridization buffer was added to cell lysate to a final concentration of 4 M urea, to denature non-specifically interacting proteins. Afterwards, the lysis of 20 M cells was pre-cleared by incubation with 125 μL of MyOne C1 Streptavidin beads (Invitrogen, USA) at 45 °C for 30 min. Biotinylated 90-mer DNA probes for RNA capture (10 μg of probes for GRAS1, 5 μg of probes each for U1 and Luciferase) were mixed with pre-cleared lysate and incubated at 45 °C for 2 hours with mixing at 1000 rpm (15 s shaking, 15 s off). Complexes of biotinylated DNA probes and RNA targets were then captured on MyOne C1 Streptavidin beads by incubating at 45 °C for 1 hour with the same mixing setting as above. Beads were washed 4 times with 4 M urea hybridization buffer, and finally resuspended with 0.5 μL of Benzonase (E1014, SIGMA-ALDRICH) in 500 μL Benzonase Elution Buffer (20 mM Tris [pH 8.0], 2 mM MgCl2, 0.05% NLS, 0.5 mM TCEP) to eluate captured proteins. Proteins were precipitated by adding 80% TCA to final concentration to 20% for overnight incubation at 4 °C. Precipitated proteins were collected by centrifugation for 15 min at 10,000 x *g* and 4 °C. The protein pellet was resuspended with 50 μL of water containing 5% Rapigest (186002123, Waters) and incubated at 100 °C for 10 min. Next, 0.5 μg of trypsin was added with 37 °C overnight incubation and 600 rpm shaking for protein digestion. Protein samples were purified using HiPPR detergent removal resin (PI88305, VWR International) and dried completely by Speedvac (Savant) before LC-MS/MS analysis at Sanford Burnham Prebys Proteomics Core on a Thermo QExactive instrument. Mass spectrometry datasets as well as MaxQuant peptide search results and parameter files are available via ProteomeXchange with identifier PXD042699.

### Annexin V apoptosis assay

Following treatment or transfection, A549 cells were collected by trypsinization and washed in cold 1X PBS. Then cells were resuspended in 100 μL of 1X Annexin-binding buffer (10 mM Hepes adjusted to pH 7.4, 140 mM NaCl and 2.5 mM CaCl2) with 5 μL of FITC-conjugated Annexin V (A13199, Life Technologies) and 2 μL of 1 mg/mL Propidium Iodide (40017, Biotium). The cell solution was incubated for 30 mins at room temperature in the dark. After the incubation, 400 μL of 1X Annexin-binding buffer was added to the cell solution and analyzed by the S3e Cell Sorter. The raw data was analyzed by FlowJo software.

## Material Availability

All unique and stable reagents generated in this study are available from the Lead Contact with a completed Materials Transfer Agreement. Further information and requests for resources and reagents should be directed to the lead contact, Colleen A. McHugh.

## Author Contributions

Conceptualization, T.S., N.T. C.A.M.; methodology, T.S., N.T., L.K., C.A.M.; validation, T.S., N.T., L.K., D.C. J.Z; formal analysis, T.S., N.T., L.K., D.C., J.Z., V.C.; investigation, T.S., N.T., L.K., L.E. D.C., J.Z., N.C., C.H., C.A.M.; supervision, C.A.M; writing and revision of draft T.S., N.T., L.K., L.E. D.C., J.Z., N.C., V.C., C.H., C.A.M.; funding acquisition, C.A.M.

## Acknowledgements

This work was supported by the Chemistry and Biochemistry Department of the University of California San Diego, the NIH/NIGMS R00 GM120494 award to C.A.M., the NIH/NCI T32 CA009523 fellowship to T.S. and N.T. (with additional support from the San Diego Fellowship), the University of California Cancer Research Coordinating Committee grant C21CR2119 to C.A.M. (with CRCC Diversity Supplement support to C.H.), the American Chemical Society Bridge Program (ACS-BP) support to N.C., and the Dean of Physical Sciences Undergraduate Summer Research Awards to L.K. and C.H. This publication includes data generated on the NovaSeq 6000 at the UC San Diego Institute for Genomic Medicine and supported by the Moores Cancer Center grant P30CA023100.

## Supplementary Figures

**Supplementary Figure 1:**
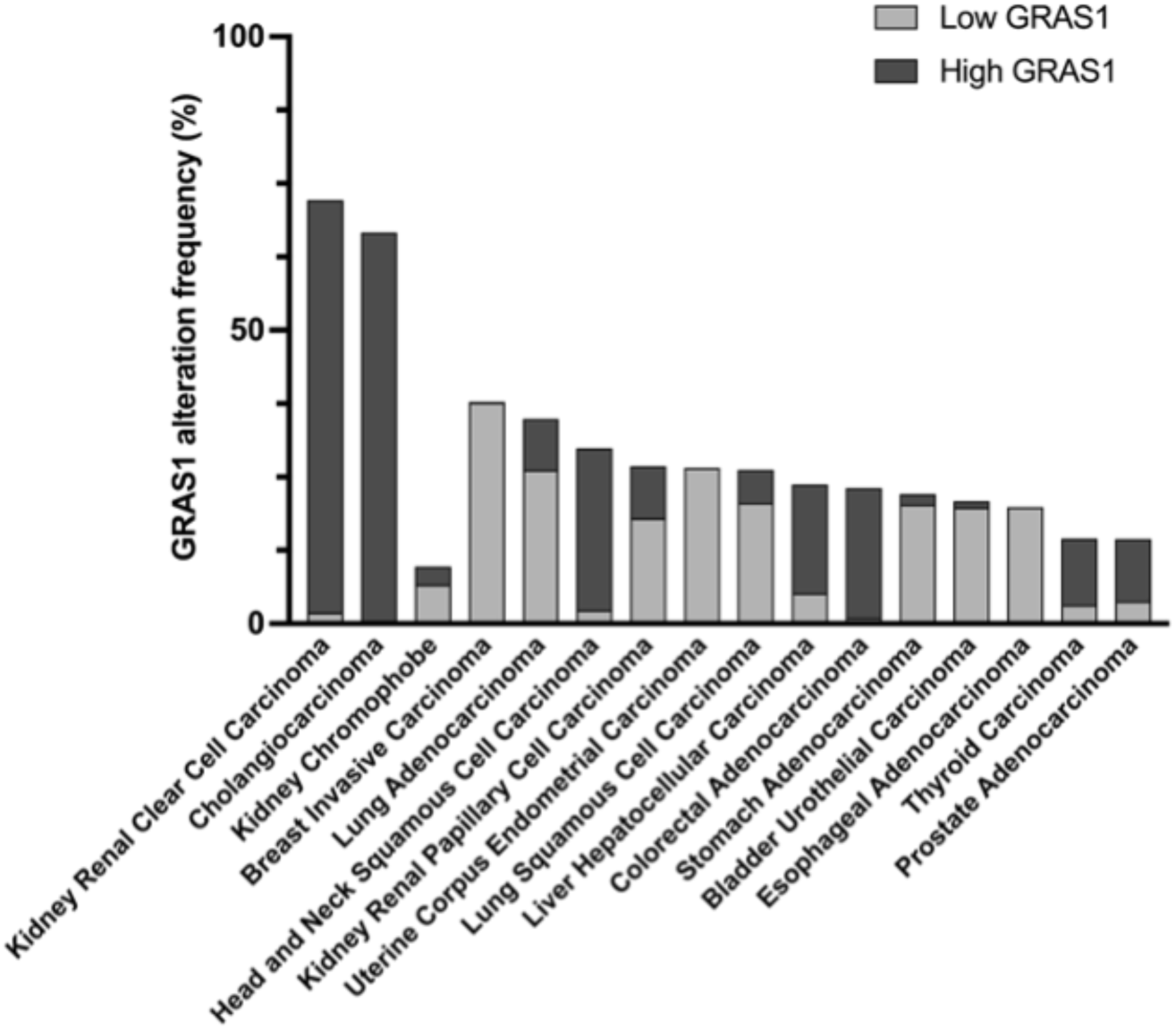
GRAS1 non-coding RNA expression is dysregulated in multiple human cancer types. TCGA PanCancer Atlas analysis of GRAS1 differential expression in tumors relative to adjacent normal tissues.

**Supplementary Figure 2:**
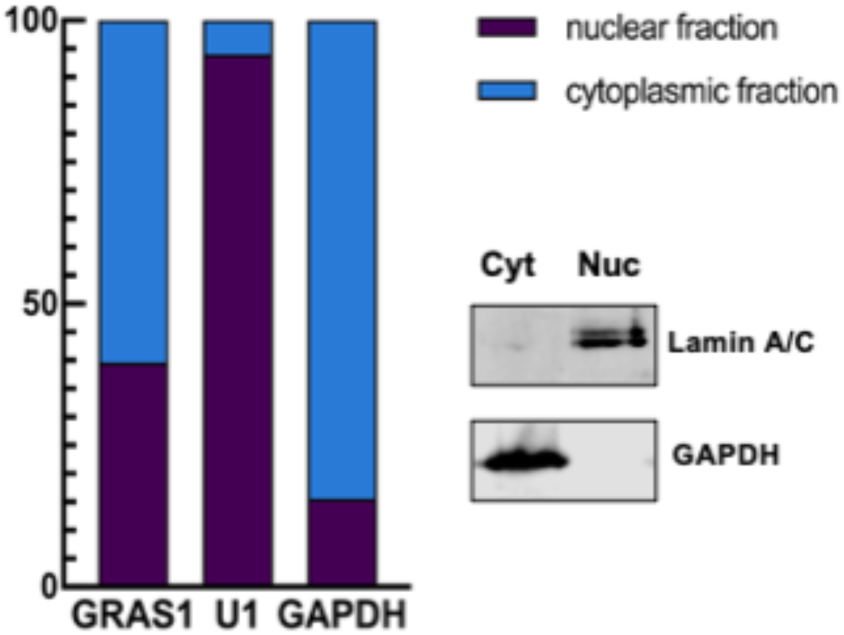
GRAS1 is localized to both the nucleus and the cytoplasm. RT-qPCR for GRAS1, U1 snRNA, and GAPDH mRNA in nuclear and cytoplasmic fractionations of A549 cells (left panel). Western blot of Lamin A/C was used as a marker for nuclear fraction, and GAPDH was used as a marker for cytoplasmic fraction (right panel).

**Supplementary Figure 3:**
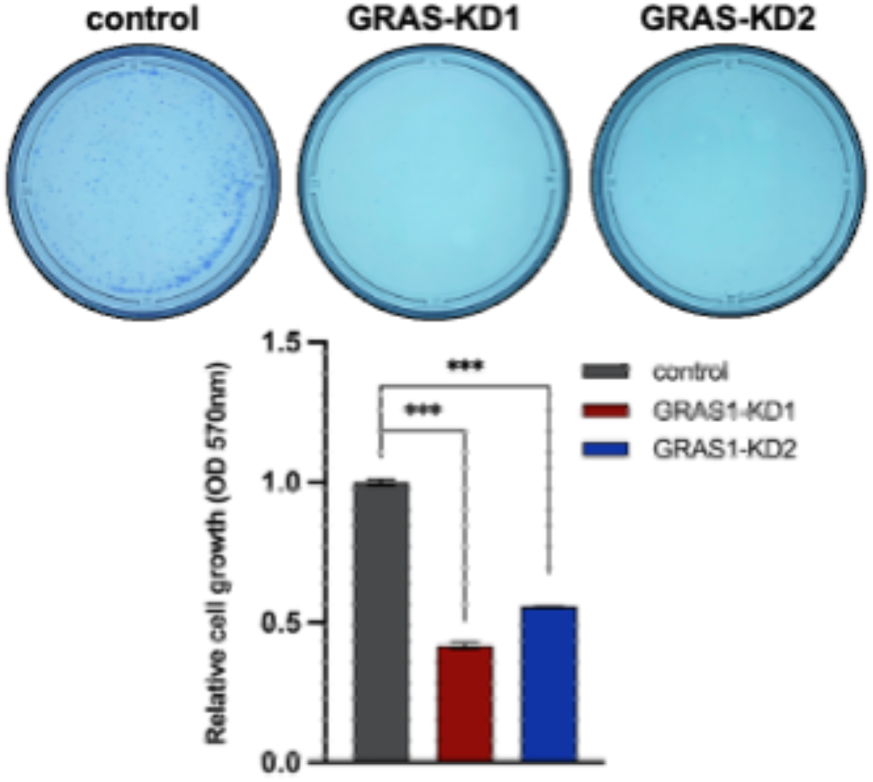
GRAS1 knockdown inhibits colony formation. Colony formation assay of A549 cells transfected with control GapmeR compared with GRAS1 GapmeR1 or GRAS1 GapmeR2. Representative cell images (top) and quantification of dissolved crystal violet staining (bottom). Error bars represent SD. (***) P < 0.001 by two-tailed Student’s t-test.

**Supplementary Figure 4:**
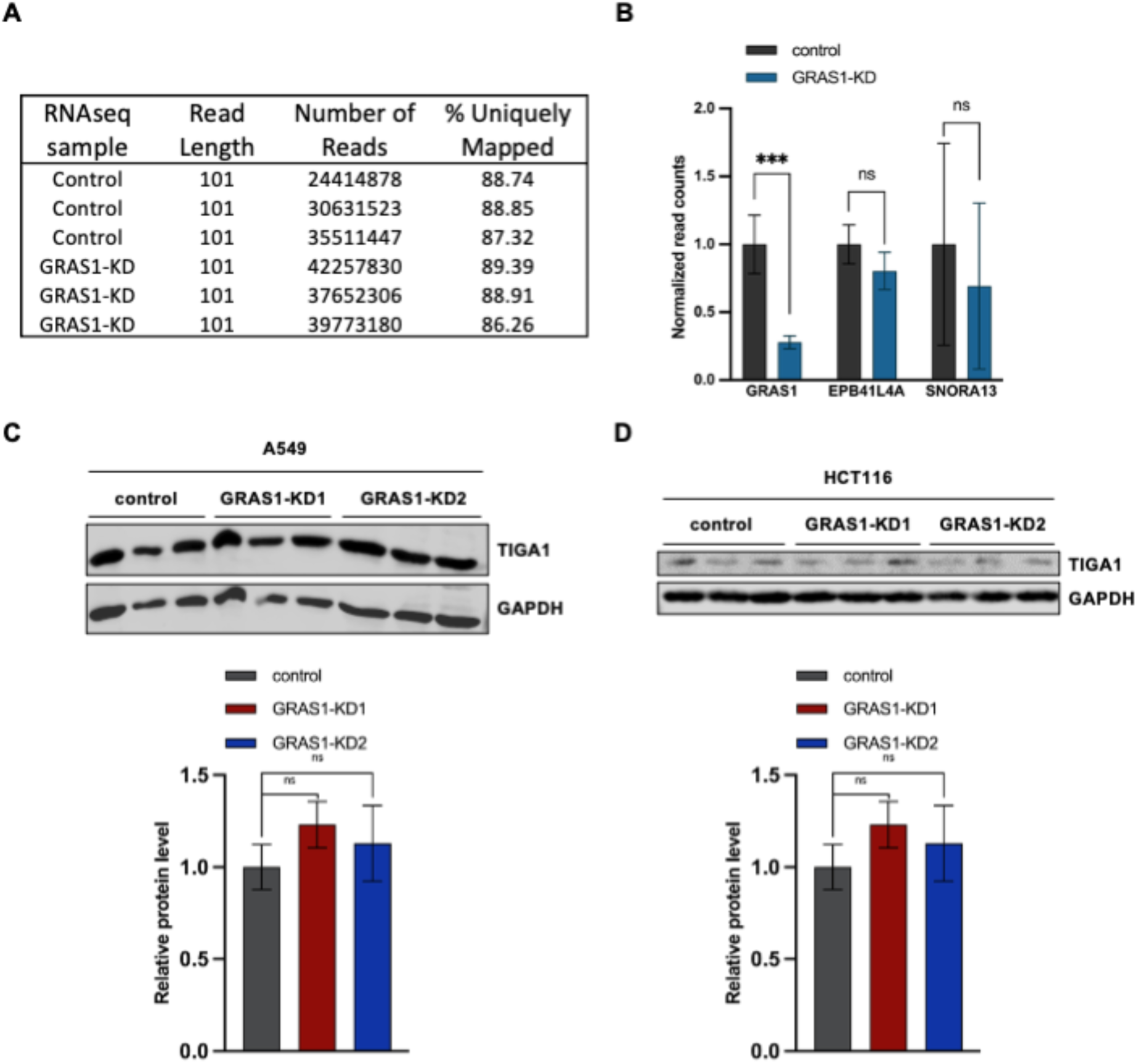
GRAS1 knockdown does not affect the expression of neighboring genes. (A) The statistics of sequence length, number of reads, and percentage of uniquely mapped in the RNA sequencing data. (B) RNA expression of GRAS1, EPB41L4A, and SNORA13 in control and GRAS1 knockdown groups based on read counts in RNA sequencing. (C) Western blot of TIGA1 in A549 cells treated with control GapmeR, GRAS1 GapmeR1 or GRAS1 GapmeR2 (top). Quantification of TIGA1 expression was calculated after normalization to the GAPDH loading control for each sample (bottom). (D) Western blot of TIGA1 in HCT116 cells treated with control GapmeR, GRAS1 GapmeR1 or GRAS1 GapmeR2 (top). Quantification of TIGA1 expression was calculated after normalization to the GAPDH loading control for each sample (bottom). Error bars represent SD. (ns) P > 0.05; (***) P < 0.001 by two-tailed Student’s t-test.

**Supplementary Figure 5:**
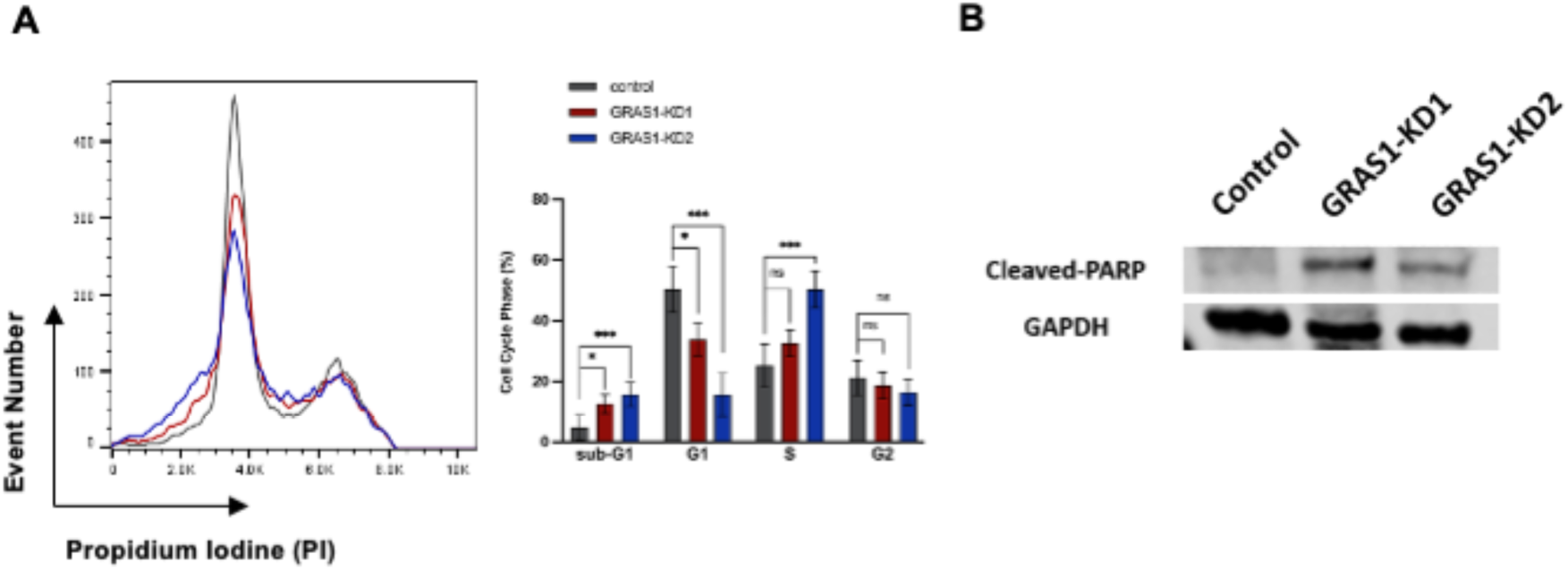
GRAS1 knockdown affects cell cycle and induces apoptosis. (A) Cell cycle analysis of GRAS1 knockdown cells. Representative images of cell cycle analysis by propidium iodide (PI) staining in flow cytometry for A549 transfected with control GapmeR, GRAS1 GapmeR1, or GRAS1 GapmeR2 (left panel). Quantification of the proportion of cells in each cell cycle phase (right panel). (B) Western blot of cleaved PARP levels in control GapmeR treated cells compared to GRAS1 GapmeR1 or GRAS1 GapmeR2 knockdown cells. Error bars represent SD. (ns) P > 0.05; (*) P ≤ 0.05; (***) P < 0.001 by two-tailed Student’s t-test.

## Supplementary Table

**Supplementary Table 1:**
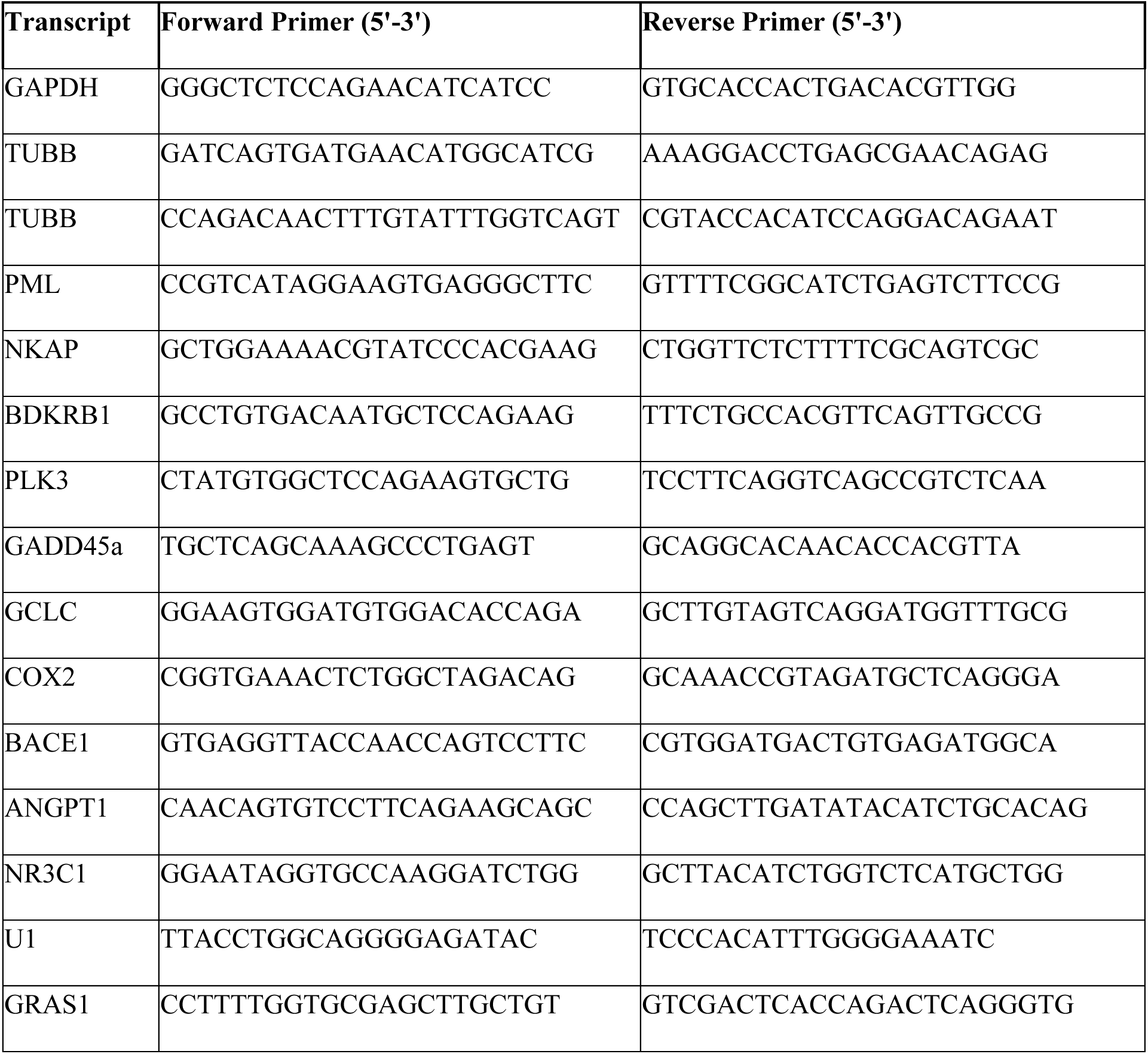
Oligonucleotide primers used for RT-qPCR analysis.

## References

Bin J, Nie S, Tang Z, Kang A, Fu Z, Hu Y, Liao Q, Xiong W, Zhou Y, Tang Y, Jiang J. Long noncoding RNA EPB41L4A-AS1 functions as an oncogene by regulating the Rho/ROCK pathway in colorectal cancer. J Cell Physiol. 2021 Jan;236(1):523–535. doi: 10.1002/jcp.29880. Epub 2020 Jun 17. PMID: 32557646.

Burgute BD, Peche VS, Steckelberg AL, Glöckner G, Gaßen B, Gehring NH, Noegel AA. NKAP is a novel RS-related protein that interacts with RNA and RNA binding proteins. Nucleic Acids Res. 2014 Mar;42(5):3177–93. doi: 10.1093/nar/gkt1311. Epub 2013 Dec 17. PMID: 24353314; PMCID: PMC3950704.

Cerami E, Gao J, Dogrusoz U, Gross BE, Sumer SO, Aksoy BA, Jacobsen A, Byrne CJ, Heuer ML, Larsson E, Antipin Y, Reva B, Goldberg AP, Sander C, Schultz N. The cBio cancer genomics portal: an open platform for exploring multidimensional cancer genomics data. Cancer Discov. 2012 May;2(5):401–4. doi: 10.1158/2159-8290.CD-12-0095. Erratum in: Cancer Discov. 2012 Oct;2(10):960. PMID: 22588877; PMCID: PMC3956037.

Chen D, Li Z, Yang Q, Zhang J, Zhai Z, Shu HB. Identification of a nuclear protein that promotes NF-κB activation. Biochem Biophys Res Commun. 2003 Oct 24;310(3):720–4. doi: 10.1016/j.bbrc.2003.09.074. PMID: 14550261.

Cui P, Zhao X, Liu J, Chen X, Gao Y, Tao K, Wang C, Zhang X. miR-146a interacting with lncRNA EPB41L4A-AS1 and lncRNA SNHG7 inhibits proliferation of bone marrow-derived mesenchymal stem cells. J Cell Physiol. 2020 Apr;235(4):3292–3308. doi: 10.1002/jcp.29217. Epub 2019 Oct 14. PMID: 31612476.

Delshad E, Shamsabadi FT, Bahramian S, Mehravar F, Maghsoudi H, Shafiee M. *In silico* identification of novel lncRNAs with a potential role in diagnosis of gastric cancer. J Biomol Struct Dyn. 2020 Apr;38(7):1954–1962. doi: 10.1080/07391102.2019.1624615. Epub 2019 Jun 8. PMID: 31179892.

Dobin A, Davis CA, Schlesinger F, Drenkow J, Zaleski C, Jha S, Batut P, Chaisson M, Gingeras TR. STAR: ultrafast universal RNA-seq aligner. Bioinformatics. 2013 Jan 1;29(1):15–21. doi: 10.1093/bioinformatics/bts635. Epub 2012 Oct 25. PMID: 23104886; PMCID: PMC3530905.

Fiordaliso SK, Iwata-Otsubo A, Ritter AL, Quesnel-Vallières M, Fujiki K, Nishi E, Hancarova M, Miyake N, Morton JEV, Lee S, Hackmann K, Bando M, Masuda K, Nakato R, Arakawa M, Bhoj E, Li D, Hakonarson H, Takeda R, Harr M, Keena B, Zackai EH, Okamoto N, Mizuno S, Ko JM, Valachova A, Prchalova D, Vlckova M, Pippucci T, Seiler C, Choi M, Matsumoto N, Di Donato N, Barash Y, Sedlacek Z, Shirahige K, Izumi K. Missense Mutations in NKAP Cause a Disorder of Transcriptional Regulation Characterized by Marfanoid Habitus and Cognitive Impairment. Am J Hum Genet. 2019 Nov 7;105(5):987–995. doi: 10.1016/j.ajhg.2019.09.009. Epub 2019 Oct 3. PMID: 31587868; PMCID: PMC6848994.

Frankish A, Diekhans M, Ferreira AM, Johnson R, Jungreis I, Loveland J, Mudge JM, Sisu C, Wright J, Armstrong J, Barnes I, Berry A, Bignell A, Carbonell Sala S, Chrast J, Cunningham F, Di Domenico T, Donaldson S, Fiddes IT, García Girón C, Gonzalez JM, Grego T, Hardy M, Hourlier T, Hunt T, Izuogu OG, Lagarde J, Martin FJ, Martínez L, Mohanan S, Muir P, Navarro FCP, Parker A, Pei B, Pozo F, Ruffier M, Schmitt BM, Stapleton E, Suner MM, Sycheva I, Uszczynska-Ratajczak B, Xu J, Yates A, Zerbino D, Zhang Y, Aken B, Choudhary JS, Gerstein M, Guigó R, Hubbard TJP, Kellis M, Paten B, Reymond A, Tress ML, Flicek P. GENCODE reference annotation for the human and mouse genomes. Nucleic Acids Res. 2019 Jan 8;47(D1):D766-D773. doi: 10.1093/nar/gky955. PMID: 30357393; PMCID: PMC6323946.

Gagnon SJ, Ennis FA, Rothman AL. Bystander target cell lysis and cytokine production by dengue virus-specific human CD4(+) cytotoxic T-lymphocyte clones. J Virol. 1999 May;73(5):3623–9. doi: 10.1128/JVI.73.5.3623-3629.1999. PMID: 10196254; PMCID: PMC104137.

Gao J, Aksoy BA, Dogrusoz U, Dresdner G, Gross B, Sumer SO, Sun Y, Jacobsen A, Sinha R, Larsson E, Cerami E, Sander C, Schultz N. Integrative analysis of complex cancer genomics and clinical profiles using the cBioPortal. Sci Signal. 2013 Apr 2;6(269):pl1. doi: 10.1126/scisignal.2004088. PMID: 23550210; PMCID: PMC4160307.

Hackmann K, Rump A, Haas SA, Lemke JR, Fryns JP, Tzschach A, Wieczorek D, Albrecht B, Kuechler A, Ripperger T, Kobelt A, Oexle K, Tinschert S, Schrock E, Kalscheuer VM, Di Donato N. Tentative clinical diagnosis of Lujan-Fryns syndrome--A conglomeration of different genetic entities? Am J Med Genet A. 2016 Jan;170A(1):94-102. doi: 10.1002/ajmg.a.37378. Epub 2015 Sep 11. PMID: 26358559.

Kastan MB, Canman CE, Leonard CJ. P53, cell cycle control and apoptosis: implications for cancer. Cancer Metastasis Rev. 1995 Mar;14(1):3–15. doi: 10.1007/BF00690207. PMID: 7606818.

Kazemzadeh M, Safaralizadeh R, Orang AV. LncRNAs: emerging players in gene regulation and disease pathogenesis. J Genet. 2015 Dec;94(4):771–84. doi: 10.1007/s12041-015-0561-6. PMID: 26690535.

Lakin ND, Jackson SP. Regulation of p53 in response to DNA damage. Oncogene. 1999 Dec 13;18(53):7644–55. doi: 10.1038/sj.onc.1203015. PMID: 10618704.

Li T, Chen L, Cheng J, Dai J, Huang Y, Zhang J, Liu Z, Li A, Li N, Wang H, Yin X, He K, Yu M, Zhou T, Zhang X, Xia Q. SUMOylated NKAP is essential for chromosome alignment by anchoring CENP-E to kinetochores. Nat Commun. 2016 Oct 3;7:12969. doi: 10.1038/ncomms12969. PMID: 27694884; PMCID: PMC5064014.

Liao Y, Smyth GK, Shi W. featureCounts: an efficient general purpose program for assigning sequence reads to genomic features. Bioinformatics. 2014 Apr 1;30(7):923–30. doi: 10.1093/bioinformatics/btt656. Epub 2013 Nov 13. PMID: 24227677.

Liu SJ, Horlbeck MA, Cho SW, Birk HS, Malatesta M, He D, Attenello FJ, Villalta JE, Cho MY, Chen Y, Mandegar MA, Olvera MP, Gilbert LA, Conklin BR, Chang HY, Weissman JS, Lim DA. CRISPRi-based genome-scale identification of functional long noncoding RNA loci in human cells. Science. 2017 Jan 6;355(6320):aah7111. doi: 10.1126/science.aah7111. Epub 2016 Dec 15. PMID: 27980086; PMCID: PMC5394926.

Liu J, Chen SJ, Hsu SW, Zhang J, Li JM, Yang DC, Gu S, Pinkerton KE, Chen CH. MARCKS cooperates with NKAP to activate NF-kB signaling in smoke-related lung cancer. Theranostics. 2021 Feb 19;11(9):4122–4136. doi: 10.7150/thno.53558. PMID: 33754052; PMCID: PMC7977464.

Love MI, Huber W, Anders S. Moderated estimation of fold change and dispersion for RNA-seq data with DESeq2. Genome Biol. 2014;15(12):550. doi: 10.1186/s13059-014-0550-8. PMID: 25516281; PMCID: PMC4302049.

Mah LJ, El-Osta A, Karagiannis TC. gammaH2AX: a sensitive molecular marker of DNA damage and repair. Leukemia. 2010 Apr;24(4):679–86. doi: 10.1038/leu.2010.6. Epub 2010 Feb 4. PMID: 20130602.

Maranon DG, Wilusz J. Mind the Gapmer: Implications of Co-transcriptional Cleavage by Antisense Oligonucleotides. Mol Cell. 2020 Mar 5;77(5):932–933. doi: 10.1016/j.molcel.2020.02.010. PMID: 32142690.

McHugh CA, Russell P, Guttman M. Methods for comprehensive experimental identification of RNA-protein interactions. Genome Biol. 2014 Jan 27;15(1):203. doi: 10.1186/gb4152. PMID: 24467948; PMCID: PMC4054858.

McHugh CA, Chen CK, Chow A, Surka CF, Tran C, McDonel P, Pandya-Jones A, Blanco M, Burghard C, Moradian A, Sweredoski MJ, Shishkin AA, Su J, Lander ES, Hess S, Plath K, Guttman M. The Xist lncRNA interacts directly with SHARP to silence transcription through HDAC3. Nature. 2015 May 14;521(7551):232-6. doi: 10.1038/nature14443. Epub 2015 Apr 27. PMID: 25915022; PMCID: PMC4516396.

Okuda H, Kiuchi H, Takao T, Miyagawa Y, Tsujimura A, Nonomura N, Miyata H, Okabe M, Ikawa M, Kawakami Y, Goshima N, Wada M, Tanaka H. A novel transcriptional factor Nkapl is a germ cell-specific suppressor of Notch signaling and is indispensable for spermatogenesis. PLoS One. 2015 Apr 14;10(4):e0124293. doi: 10.1371/journal.pone.0124293. PMID: 25875095; PMCID: PMC4397068.

Pajerowski AG, Nguyen C, Aghajanian H, Shapiro MJ, Shapiro VS. NKAP is a transcriptional repressor of notch signaling and is required for T cell development. Immunity. 2009 May;30(5):696–707. doi: 10.1016/j.immuni.2009.02.011. Epub 2009 Apr 30. PMID: 19409814; PMCID: PMC2777751.

Roberts TC, Morris KV, Weinberg MS. Perspectives on the mechanism of transcriptional regulation by long non-coding RNAs. Epigenetics. 2014 Jan;9(1):13–20. doi: 10.4161/epi.26700. Epub 2013 Oct 22. PMID: 24149621; PMCID: PMC3928176.

Schneider VA, Graves-Lindsay T, Howe K, Bouk N, Chen HC, Kitts PA, Murphy TD, Pruitt KD, Thibaud-Nissen F, Albracht D, Fulton RS, Kremitzki M, Magrini V, Markovic C, McGrath S, Steinberg KM, Auger K, Chow W, Collins J, Harden G, Hubbard T, Pelan S, Simpson JT, Threadgold G, Torrance J, Wood JM, Clarke L, Koren S, Boitano M, Peluso P, Li H, Chin CS, Phillippy AM, Durbin R, Wilson RK, Flicek P, Eichler EE, Church DM. Evaluation of GRCh38 and de novo haploid genome assemblies demonstrates the enduring quality of the reference assembly. Genome Res. 2017 May;27(5):849–864. doi: 10.1101/gr.213611.116. Epub 2017 Apr 10. PMID: 28396521; PMCID: PMC5411779.

Shapiro MJ, Anderson J, Lehrke MJ, Chen M, Nelson Holte M, Shapiro VS. NKAP Regulates Senescence and Cell Death Pathways in Hematopoietic Progenitors. Front Cell Dev Biol. 2019 Oct 2;7:214. doi: 10.3389/fcell.2019.00214. PMID: 31632967; PMCID: PMC6783958.

Shu W, Liu G, Dai Y, Feng A, Chen Z, Han J, Li X. The oncogenic role of NKAP in the growth and invasion of colon cancer cells. Oncol Rep. 2019 Nov;42(5):2130–2138. doi: 10.3892/or.2019.7322. Epub 2019 Sep 18. PMID: 31545474.

Shu CH, Yang WK, Shih YL, Kuo ML, Huang TS. Cell cycle G2/M arrest and activation of cyclin-dependent kinases associated with low-dose paclitaxel-induced sub-G1 apoptosis. Apoptosis. 1997;2(5):463–70. doi: 10.1023/a:1026422111457. PMID: 14646529.

Sun S, Gao T, Pang B, Su X, Guo C, Zhang R, Pang Q. RNA binding protein NKAP protects glioblastoma cells from ferroptosis by promoting SLC7A11 mRNA splicing in an m^6^A-dependent manner. Cell Death Dis. 2022 Jan 21;13(1):73. doi: 10.1038/s41419-022-04524-2. PMID: 35064112; PMCID: PMC8783023.

Toné S, Sugimoto K, Tanda K, Suda T, Uehira K, Kanouchi H, Samejima K, Minatogawa Y, Earnshaw WC. Three distinct stages of apoptotic nuclear condensation revealed by time-lapse imaging, biochemical and electron microscopy analysis of cell-free apoptosis. Exp Cell Res. 2007 Oct 1;313(16):3635–44. doi: 10.1016/j.yexcr.2007.06.018. Epub 2007 Jul 4. PMID: 17643424; PMCID: PMC2705844.

Trang N, Su T, Hall S, Boutros N, Kong B, Huang C, McHugh CA. Streamlined Purification of RNA-Protein Complexes Using UV Cross-Linking and RNA Antisense Purification. Methods Mol Biol. 2023;2666:213–229. doi: 10.1007/978-1-0716-3191-1_16. PMID: 37166668.

Uramoto H, Tanaka F. Recurrence after surgery in patients with NSCLC. Transl Lung Cancer Res. 2014 Aug;3(4):242–9. doi: 10.3978/j.issn.2218-6751.2013.12.05. PMID: 25806307; PMCID: PMC4367696.

Wang M, Zheng S, Li X, Ding Y, Zhang M, Lin L, Xu H, Cheng Y, Zhang X, Xu H, Li S. Integrated Analysis of lncRNA-miRNA-mRNA ceRNA Network Identified lncRNA EPB41L4A-AS1 as a Potential Biomarker in Non-small Cell Lung Cancer. Front Genet. 2020 Sep 18;11:511676. doi: 10.3389/fgene.2020.511676. PMID: 33193600; PMCID: PMC7530329.

Wei Y, Wang Y, Zang A, Wang Z, Fang G, Hong D. MiR-4766-5p Inhibits The Development And Progression Of Gastric Cancer By Targeting NKAP. Onco Targets Ther. 2019 Oct 16;12:8525-8536. doi: 10.2147/OTT.S220234. PMID: 31802890; PMCID: PMC6801498.

Yang F, Lv S. LncRNA EPB41L4A-AS1 Regulates Cell Proliferation, Apoptosis and Metastasis in Breast Cancer. Ann Clin Lab Sci. 2022 Jan;52(1):3–11. Erratum in: Ann Clin Lab Sci. 2022 May;52(3):510. PMID: 35181612.

Yu G, Wang LG, Han Y, He QY. clusterProfiler: an R package for comparing biological themes among gene clusters. OMICS. 2012 May;16(5):284–7. doi: 10.1089/omi.2011.0118. Epub 2012 Mar 28. PMID: 22455463; PMCID: PMC3339379.

Zhang X, Duan J, Li Y, Jin X, Wu C, Yang X, Lu W, Ge W. NKAP acts with HDAC3 to prevent R-loop associated genome instability. Cell Death Differ. 2023 Jul;30(7):1811–1828. doi: 10.1038/s41418-023-01182-5. Epub 2023 Jun 15. PMID: 37322264; PMCID: PMC10307950.

Zhu Y, Liu Q, Liao M, Diao L, Wu T, Liao W, Wang Z, Li B, Zhang S, Wang S, Xie W, Jiang Y, Xu N, Zeng Y, Yang BB, Zhang Y. Overexpression of lncRNA EPB41L4A-AS1 Induces Metabolic Reprogramming in Trophoblast Cells and Placenta Tissue of Miscarriage. Mol Ther Nucleic Acids. 2019 Dec 6;18:518–532. doi: 10.1016/j.omtn.2019.09.017. Epub 2019 Sep 26. PMID: 31671345; PMCID: PMC6838551.

